# Genome-wide cline analysis identifies new locus contributing to a barrier to gene flow across an *Antirrhinum* hybrid zone

**DOI:** 10.1101/2025.02.17.638607

**Authors:** David L Field, Sean Stankowski, Taylor Reiter, Jitka Polechova, Desmond Bradley, Daniel Richardson, Arka Pal, Daria Shipilina, Louis Boell, Melinda Pickup, Yongbiao Xue, Enrico Coen, Nicholas Barton

## Abstract

Identification of the genomic regions that contribute to reproductive isolation and how they interact is a major goal of evolutionary genetics. Much effort has focused on locating candidate genes and potential barrier loci by scanning genomes for regions of excess differentiation (*F_ST_*). An alternative, and perhaps more robust approach, would be to scan for genomic regions exhibiting steep clines in allele frequency across a hybrid zone. Here we develop this approach and apply it to genomic data from across a hybrid zone between varieties of *Antirrhinum majus* subspecies *majus* with different flower colour patterns (*A. m. m* var. *pseudomajus* and *A. m. m* var. *striatum*). Most steep clines are clustered in seven genomic regions, only four of which were present from *F_ST_* scans between all pair-wise comparisons. Six of these regions carry previously identified loci that influence flower colour in the hybrid zone. The seventh region harbours a novel locus*, RUBIA*, modifying magenta intensity. Clines at *RUBIA* approached towards fixation on magenta side of the hybrid zone, whilst remaining polymorphic on the yellow side. This polymorphism on the yellow side may reflect a smaller phenotypic effect of *RUBIA* in yellow compared to magenta genetic backgrounds. Our findings illustrate how whole-genome cline scans in hybrid zones can robustly detect genomic regions contributing to phenotypic differences and highlight how different reproductive barrier loci interact across the genome.

## Introduction

A fundamental question in evolutionary biology is how divergent populations arise and are maintained in the face of gene flow (Ravinet et al., 2017; Wolf & Ellegren, 2017). The separation of species ultimately requires the evolution of barriers to gene exchange that are strong enough to balance the homogenizing effects of gene flow. Much effort has focused on identifying the genetic barriers to gene flow and the traits underlying reproductive isolation (Ravinet et al., 2017). The heterogenous landscape of phylogenetic relationships (e.g. Martin et al., 2019) and genome-wide scans of population differentiation and divergence (*F_ST_* and *D*_xy_) have been widely used strategies for locating barrier loci along genomes (Wolf & Ellegren, 2017). The *F_ST_* scan approach assumes that regions of excess divergence (genomic islands) coincide with barrier loci whereas regions of low divergence reflect homogenization through recurrent gene flow (Feder et al., 2012; Wu, 2001). However, patterns of differentiation are often complex and genomic islands may be produced by a number of processes unrelated to differential gene flow (Bierne, 2010; Cruickshank & Hahn, 2014a; Wolf & Ellegren, 2017; Wong & Filatov, 2023). Although useful insights have been developed to account for confounding effects of background selection and recombination (e.g. Martin et al., 2019; Stankowski et al., 2019), these signals still often tell us little about the ongoing balance between gene flow and selection required to demonstrate local barriers along the genome.

Genome-wide scans for regions exhibiting geographic clines in allele frequency across hybrid zones provide a promising way to find genomic regions containing barriers to gene flow. In hybrid zones, genomes continually mix to produce new gene combinations which selection acts upon in nature. If these are less fit, either inherently or because they are in an unfavorable environment, then a stable equilibrium may be reached between gene flow (dispersal) and selection (Haldane, 1948; Slatkin, 1973; Barton & Hewitt, 1985). This leaves a signature of steep geographic clines in allele frequency, which can be quantified for each locus along the genome. Theory predicts that divergently selected alleles are expected to resist introgression and maintain steep clines, whereas neutral or advantageous alleles will exchange freely across the hybrid zone and eventually flatten out. A key strength of geographic cline analysis is that local equilibrium is reached quickly for selected (*t*=1/*s*) and neutral loci (*t*=x^2^/*σ*^!^for distance x) (Barton & Gale, 1993; Barton & Hewitt, 1985). Therefore, given enough generations since secondary contact, loci under divergent selection can be located along the genome because they display steeper spatial gradients in allele frequency compared to neutral loci.

There is a long history of fitting geographic clines to small numbers of loci to estimate selection in hybrid zones (Endler, 1977; Haldane, 1948) with increasing numbers of loci available as sequencing technologies improved (e.g. Carneiro et al., 2013; Macholán et al., 2011). Although a few studies have begun scaling up to many thousands of loci, most still use specific loci of interest or sparse reduced representation genomic data (Stankowski et al., 2023; Westram et al., 2018). As the field moves toward whole genome resequencing data (Rafati et al., 2018), the increasing computational burden of fitting complex cline models to large numbers of loci requires the development of new tools for efficient genome wide cline estimation. This would allow for rapid un-ascertained scanning of genome wide clines, their distribution and parameters, which would be valuable for identifying barrier loci and genes responsible for divergent phenotypic traits.

Shifting to whole genomic scans of clines raises new statistical challenges. Most traditional cline fitting procedures are based on comparing observed frequencies to expected frequencies across a range of cline models using Maximum Likelihood Estimation (MLE) assessed with simulated annealing or Metropolis-Hastings algorithms (e.g. Barton & Baird, 1995; Derryberry et al., 2014). For large numbers of loci this is a considerable computational burden, and so the development of a faster routine as a complementary method to traditional fitting would be a valuable tool for studying hybrid zones. Once cline properties are estimated genome wide, it is unclear how they will be distributed across genomes in relation to known positions of selected loci. Furthermore, it remains unclear whether selected loci can be reliably picked out against the stochasticity that we expect across large numbers of loci. Recent attempts to simulate genome wide geographic clines have been based on simulating selected and neutral loci to provide expectations for observed cline properties (e.g. Westram et al., 2018). However, the distribution of genome wide cline properties will depend on several parameters that may be difficult to measure in nature, including: the time since secondary contact, neighbourhood size, genome wide barriers, epistasis and local recombination rate. Thus, interpreting patterns of genome wide clines is still challenging, knowing at least some true positives (i.e. genes with known phenotypic effects under divergent selection) allows inferences to be validated and provides an important reference to compare against the genomic background.

In this study, we developed a new method for geographic cline fitting and use it to locate a novel barrier locus in a hybrid zone between varieties of snapdragon *Antirrhinum majus* subspecies *majus*. This subspecies has been a model plant since the dawn of genetics (Schwarz-Sommer et al., 2003). Here, we focus on a hybrid zone between two varieties of *A. majus* subspecies *majus*. *Antirrhinum majus majus* var. *pseudomajus* is predominately magenta, in contrast with *A. m. m.* var. *striatum,* which is predominantly yellow (Fig 1). These phenotypes are determined by the interaction of a few, large effect loci, that regulate different components of the flavanol biosynthetic pathway and influence the pattern of pigmentation of anthocyanin (magenta) and aurone (yellow). The magenta colour is primarily controlled by two tightly linked MYB-like transcription factors on chromosome 6, *ROSEA* (*ROS*) (Schwinn et al., 2006; Whibley et al., 2006) and *ELUTA* (*EL*) (Tavares et al., 2018). Yellow colour is controlled by *SULFUREA* (*SULF*) on chromosome 4 (Bradley et al., 2017), *FLAVIA* (*FLA*) and *AURINA* (*AUN*) on chromosome 2 (Bradley et al., in prep), and *CREMOSA* (*CRE*) on chromosome 1 (Richardson et al., in prep). This set of flower colour genes, all of which have been shown to likely act as barriers to gene flow, makes snapdragons a useful yardstick to compare clines at barrier loci against the rest of the genome.

**Figure 1.**
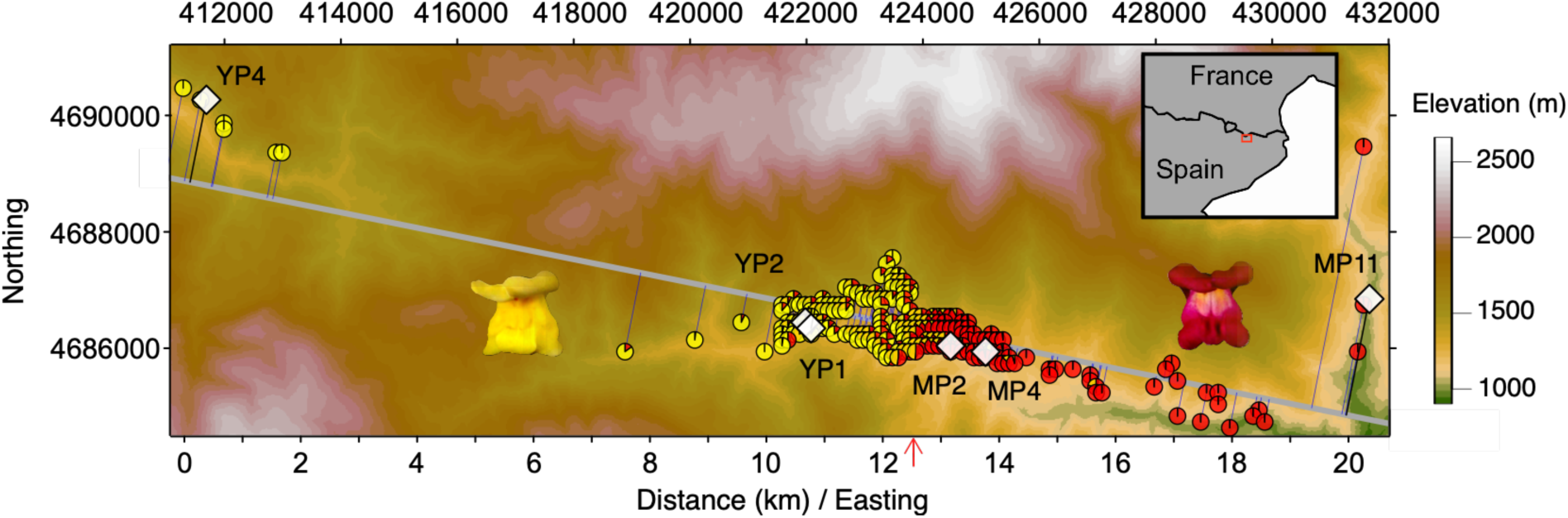
Sampling locations and allele frequencies for *Antirrhinum majus majus* hybrid zone that occurs from varieties *A. m. m.* var. *striatum* (yellow) and *A. m. m.* var. *pseudomajus* (magenta) in the Spanish Pyrenees. Sampling locations of six poolSeq whole genomes (white diamonds) and allele frequencies at *ROS* from denser sampling of individuals where each pie diagram shows the proportion of the *ROS^ps^* allele from *pseudomajus* (red) compared to the *ros^st^* allele from *striatum* (yellow) alleles at the *ROS* locus within a 200 metre diameter (see methods). Red arrow indicates the approximate centre of the phenotype cline.

The two colour varieties of *Antirrhinum majus* subspecies *majus* likely diverged over many millennia, through drift and selection and came into contact recently following glacial retreat in northern Spain. Previous studies found no evidence for post-zygotic barriers (Andalo et al., 2010), yet steep clines in colour phenotype and allele frequencies at *ROS* (Whibley et al., 2006), *EL* (Tavares et al., 2018) and *SULF* (Bradley et al., 2017), is consistent with selection acting on genes responsible for magenta and yellow pigmentation. In some cases, these regions are punctuated by narrow genomic islands of differentiation (Tavares et al., 2018). Whether steep clines are restricted only to regions with known genes that influence flower colour, cluster near genomic islands, or are dispersed throughout the genome remains unclear. How clines are distributed and their properties in relation to these features, would be useful for understanding how well different approaches identify barriers and additional genes responsible for colour variation.

Using six whole genome pools spanning the hybrid zone, we ask whether genome-wide geographic clines are steeper at loci known to influence flower colour in the hybrid zone. We also ask whether these clines are clustered together along the genome, associated with genomic islands of differentiation or with known colour loci. With whole genome pools at different distances across a transect through the hybrid zone, we expect differences between the most distant pools to reflect ancient divergence. In contrast, sharp divergence between adjacent samples may reflect selection and barriers to gene flow, maintaining divergence and steep clines at particular positions along the genome. Using a novel method for cline parameter estimation, we show that most genomic regions with steep clines are tightly clustered in genomic regions containing previously identified flower colour loci. Clines are also tightly clustered in only a few regions of the genome. By interrogating a new region enriched for clinal SNPs, we use genotypes and colour phenotype associations together with RNAseq, to identify a previously undescribed colour locus. This new gene, *RUBIA* (*RUB*) modifies the intensity of magenta colouration of the flower. Clines at this locus are asymmetric, with the cline going close to fixation on the magenta side of the hybrid zone yet polymorphic on the yellow side. This asymmetry is consistent with *RUB* having a lesser phenotypic effect in a predominantly yellow background and thus is likely under weaker selection on the yellow side of the hybrid zone. By examining genome wide properties of geographic clines, our study provides a novel contribution to locating barrier loci in relation to genomic divergence and how multiple genes interact to generate phenotypic variation under selection in nature.

## Results

### Genome wide pools along the hybrid zone highlight few fixed differences

To investigate genome-wide geographic clines and measures of nucleotide diversity (*π*), differentiation (*F_ST_*) and divergence (*D*_xy_), we used poolSeq to obtain whole-genome allele-frequency data from six sample locations spanning the transition in flower colour across the hybrid zone. The six demes were arrayed along a 1-D transect spanning 24 km (see Tavares et al., 2018), with tighter spacing of demes near the centre of the phenotypic cline to capture the steep transition (Fig 1; locations see Table S1).

The sequence data covered 476.63 Mbp (93.6%) of the ∼509 Mbp *Antirrhinum majus* reference genome v3.5 (Li et al., 2019; Zhu et al., 2023). After filtering (low sequencing depth, singletons and non-biallelic SNPs), the median of 79.7% of sites within the 10,000 bp sliding windows was covered with 46,100 windows (93.7%) remaining with the minimum of 10% window coverage. Average depth within windows ranged from 22.5 for pool MP2 to 53.3 for pool YP2. Along the genome, we identified ∼2.1 x 10^7^ variant sites across the hybrid zone.

Allele frequencies were generally similar between the two variants of *A. m. majus*, with only 3,847 (0.018% of variant sites) exhibiting allele frequency differences (Δ*p*) greater than 0.9 between the most geographically distant pools (YP4 and MP11) and 2,319 sites (0.011% of variant sites) exhibiting fixed differences (Δ*p* = 1). These fixed differences were found across the genome, although chromosome 2 contained the majority with 1,995 (86%), followed by chromosome 6 with 188 (5.1%). The remaining chromosomes exhibited much lower numbers of fixed differences (between 2 on Chr 7 and 73 on Chr 4).

### Steep clines and discordant cline centres around known flower colour loci

To facilitate the estimation of geographic cline parameters for whole-genome data, we developed a new method for rapid approximation of cline properties. The challenge with the most commonly used MLE method is they are computationally demanding for whole genome data. A simple surrogate is to estimate cline width via total heterozygosity and cline centre via centre mass of allele frequencies along a 1-dimensional transect (Polechová & Barton, 2011). We extend this approach in a method we call *FastClines*, to account for non-diagnostic loci and uneven positioning of demes (Fig 2, see methods). We use *FastClines* at 3,847 highly divergent loci (Δ*p* ≥ 0.90 between most distant pools YP4 and MP11) and use cline width to estimate the steepness of the transition and centre to estimate cline position along the transect. Cline widths ranged from ∼12.5km to 0.8km, narrowing as the cline centres moved either side of the hybrid zone (< 8km and >15km) (Fig 3). This narrowing of cline widths towards the edges is consistent with simulations of clines with six demes and reflects a limitation of small numbers of demes (Fig S1). Therefore, in the current *Antirrhinum* data set, we cannot accurately compare the steepness of clines that are shifted far to the left or right and restrict subsequent analyses to clines centred between 9km and 15km (see Fig 3). Despite this artefact, simulations of clines that incorporate sources of error (sampling and sequencing) show that different cline widths can be distinguished even with only six demes across a hybrid zone with fixed differences with power increasing with the number of demes (text S2, Fig S1). Cline properties can be distinguished as loci become less differentiated on either side of the cline although decreasing sequencing depth, variance in allele frequencies makes these estimates coarser (Fig S2).

**Figure 2.**
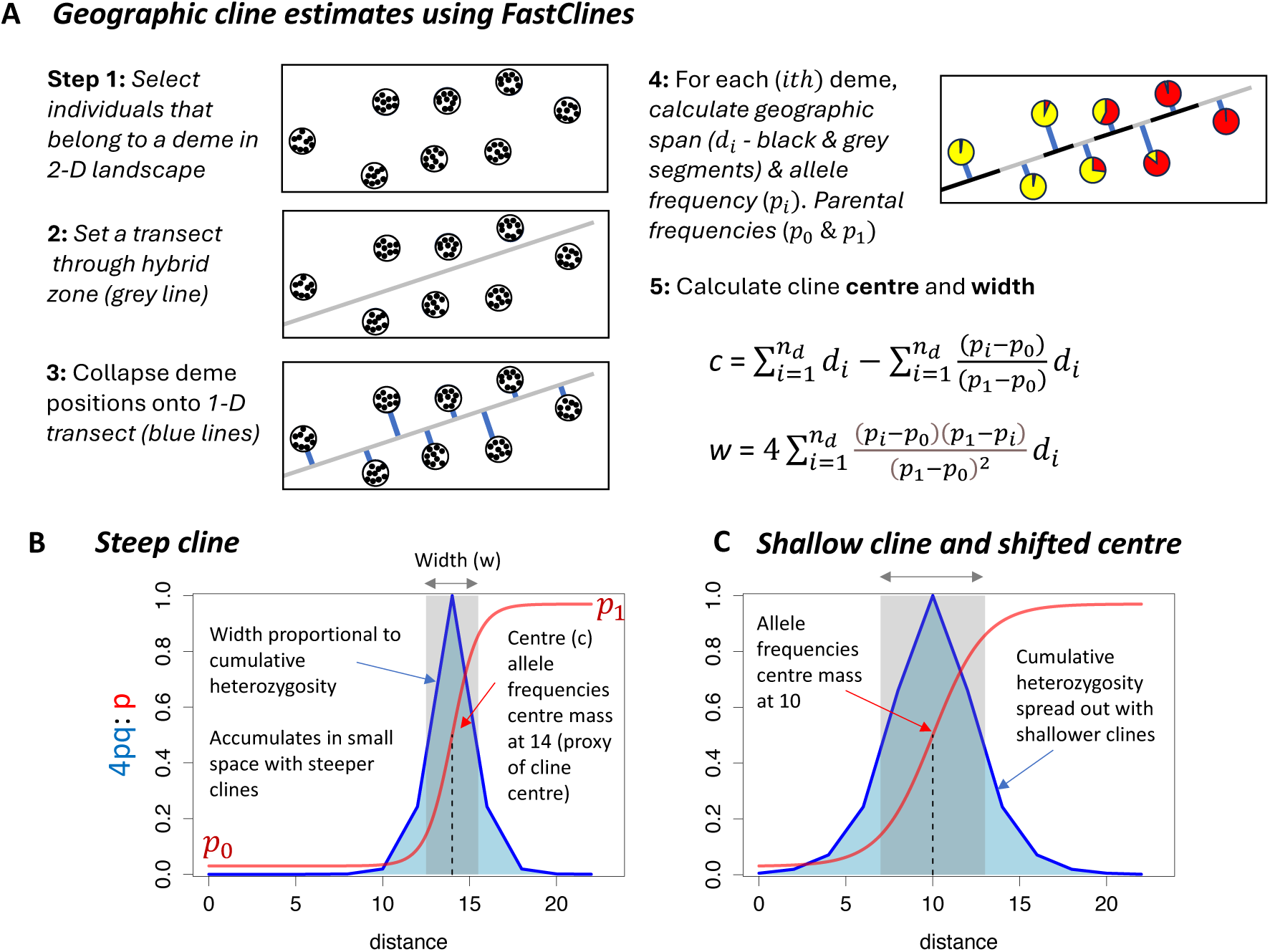
Summary of geographic clines estimation using FastClines. **(A)** Steps involved in estimating cline properties (centre and width) using FastClines with genotype data from natural populations across a hybrid zone. **(B)** cumulative heterozygosity (*4pq*) and allele frequencies (*p*) for a theoretical steep cline (width = 3, centre = 14) and the effects on the total heterozygosity across the transect (blue shaded area), **(C)** a theoretical shallower cline with a shifted centre (width = 6, centre = 10) displaying a broader spread of heterozygosity.

**Figure 3.**
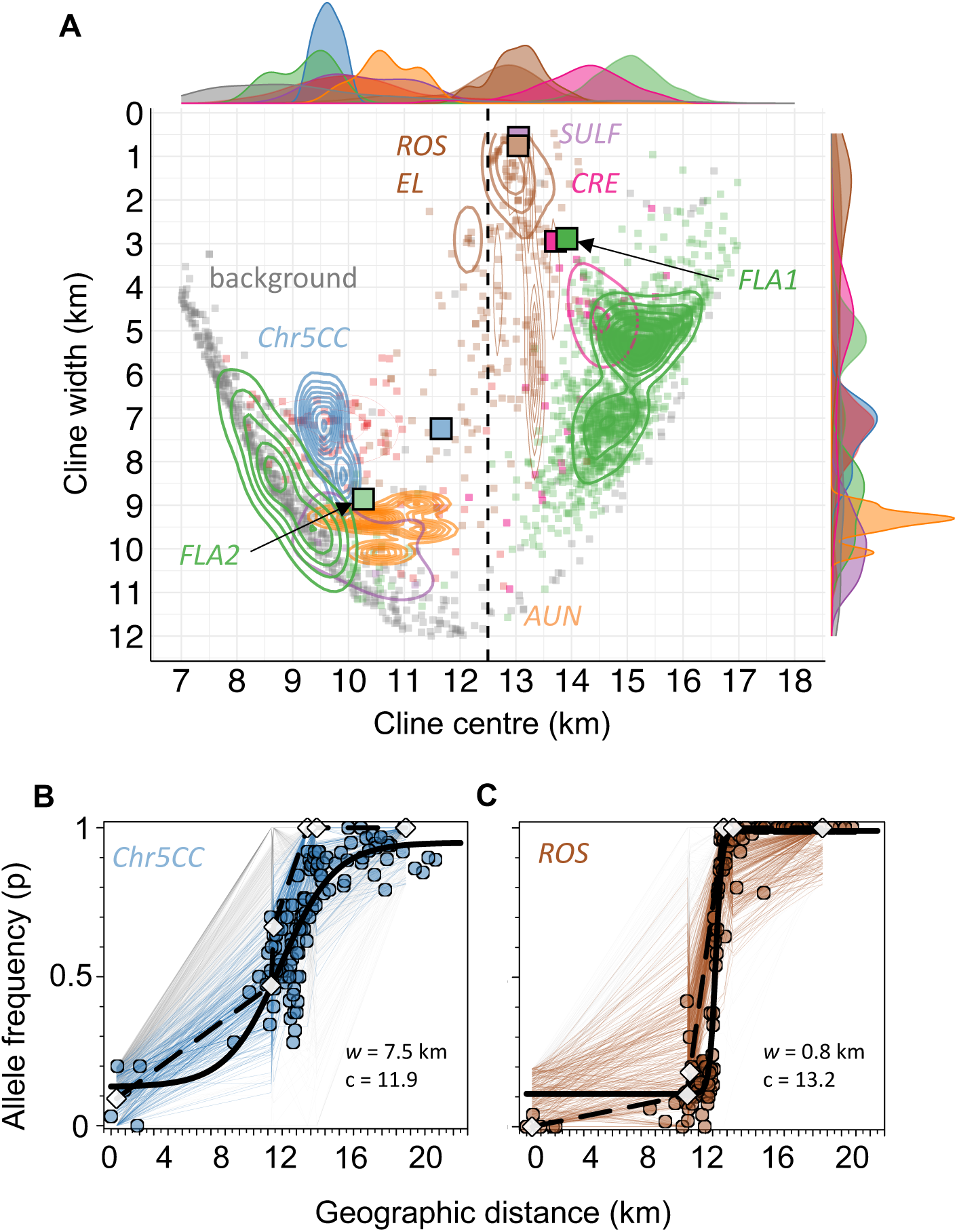
Geographic cline properties at the *Antirrhinum majus majus* hybrid zone with poolSeq estimates with FastClines and traditional cline fitting with SNP genotyping. **(A)** FastClines estimates of cline width and centre using poolSeq (circles) and traditional Maximum Likelihood Estimates (MLE) using SNP genotyping (squares). For poolSeq data using six demes, loci linked (<100kb) to colour genes (*ROS & EL* = brown, *CRE* = pink, Chr5CC/RUB = blue, *FLA1* = darker green, *FLA2* = light green, *SULF* = purple, *AUN* = orange) and background loci unlinked to colour loci (grey circles). *FLA1* refers to the cluster of loci on one side of the gene situated > 13km centre positions, and *FLA2* refers to the cluster of loci on the other side of the gene with < 13km centre positions. Contours indicate density of cline property estimates. The MLE cline parameter estimates at the four representative loci from the same regions using dense geographic sampling (>100 demes). Marginal curves indicate density of cline properties for *FastClines* (the same colours as described above). **(B)** Chromosome 5 allele frequencies from poolSeq at a locus within *RUB* (white diamond and dashed black line) for loci tightly linked to Chr5CC/*RUB* (blue lines, <100kb), background loci (grey lines, >100kb), and SNP genotyping for same locus within Chr5CC*/RUB* (blue circles) with MLE cline fit (solid black curve). **(C)** Chromosome 6 allele frequencies from poolSeq at a locus within *ROS* (white diamond and dashed black line) for loci tightly linked to *ROS1* (brown lines, <100kb), background loci (grey lines, >100kb), and SNP genotyping for same locus within Chr5CC*/RUB* (brown circles) with MLE cline fit (solid black curve).

We then mapped the genomic location of all genes known to influence flower colour to compare cline properties. The steepest clines overall were observed for loci tightly linked to *ROS* and *EL*, the two major effect loci important for regulating magenta pigmentation (Fig 3; Fig 4). For 391 clinal loci centred closer to the core region near the phenotypic centre for colour (between 11 and 14 km), the steepest clines were for loci tightly linked to *ROS* and *EL*, the two major effect loci important for regulating magenta pigmentation (Fig 3b). Of the clines around *ROS* and *EL*, clines were steepest within coding sequences for the MYB genes with mean cline width of 2.2km (±1.2km SD) at *ROS1*, 1.3km (±0.3km SD) and 3.5km (±2.3km SD) at *EL* (Fig 4a). In this region, clinal loci tended to become wider as loci were located further away from the coding sequences of these two genes. For example, 3.9km mean width for loci <30kb from *EL*, increased to 5.8km for loci 30 to 100kb away (Fig 4a).

**Figure 4.**
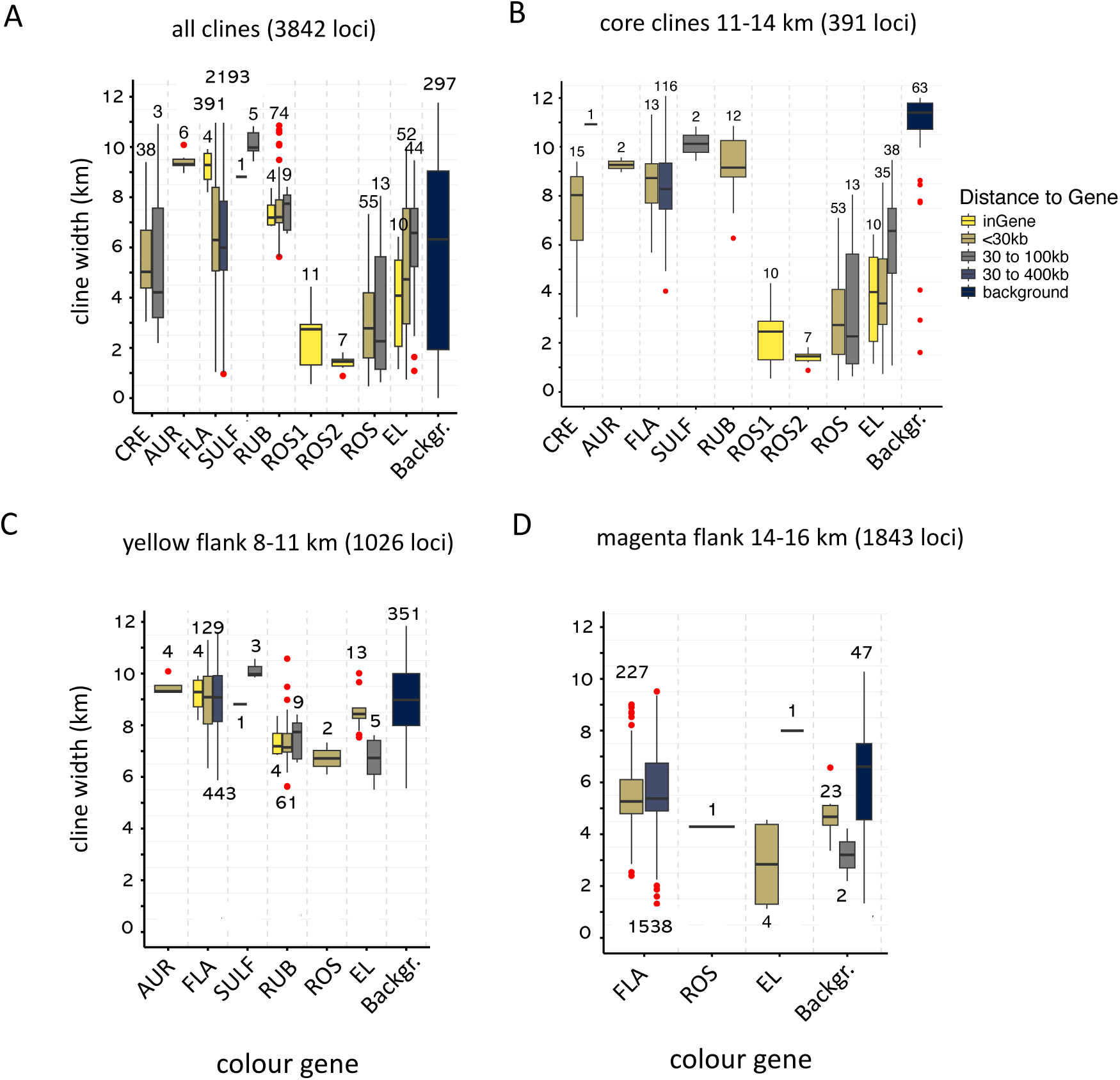
Summary of cline numbers and cline widths at different distances for colour loci for **(A)** all clines, **(B)** clines centred in the core region close to phenotypic transition, **(C)** clines centred towards the yellow flank of hybrid zone, **(D)** clines centred towards the magenta flank of the hybrid zone. Clinal loci grouped by proximity to colour loci including within the coding sequence of the gene (inGene), tightly linked < than 30kb, either 30 to 400 kb for *FLA* gene due to low recombination region, 30 to 100 kb (for all other colour loci) and background loci (>400kb near *FLA* region and >100kb elsewhere). *ROS1* and *ROS2* are loci within MYB genes, whereas *ROS* and *EL* are linked (<30kb) yet outside the coding regions of MYBs.

The ability of steep clines using *FastClines* to detect other major colour loci depended on where the cline centre was located. Cline centre displayed a range of positions either side of the main phenotypic transition. A large number of loci (n = 1843) displayed a cline centre to the east of the phenotype centre (magenta side). The discordance in cline centre among loci was strongly associated with genomic location, with 1765 of these 1843 loci (95%) with cline centre >14 km, occurring on chromosome 2 immediately downstream of the *FLA* gene (see *FLA1* for downstream sites on Fig 3a). In contrast, clinal loci within the coding sequence of *FLA* and upstream show a change in clines centre < 13 km (see *FLA2* for upstream sites on Fig3a; see Fig S5). The majority of ‘background’ loci which were greater than 300kb distant from known colour loci exhibited cline centres shifted into the yellow side (grey points; Fig 3a). These loci showed a typical pattern of high frequencies of the common allele from the magenta flanks in five of the six pools, followed by a rapid drop in allele frequency in the YP4 pool located on the other side of the mountain pass. This pattern of allele frequency change was evident for loci not closely linked to colour genes (>300kb *FLA* on Chr 2 see Fig S6; >100kb from *CRE* on Chr 1 see Fig S7, >100kb from *SULF* on Chr 4 see Fig S8; >100kb from Chr5CC see Fig S9; >100kb from *ROS*/*EL* on Chr 6 see Fig S10) and for most ‘background’ loci on chromosomes which do not possess any known colour loci (Chr 3, Chr 7 and Chr 8 see Fig S11).

Next we focus on loci with clines positioned closer to the phenotype transition between 8-16km. Given the tendency for widths to narrow as cline position shifts to the edge of the transect, together with the sharp allele frequency step for loci positioned over the mountain pass, this ensures we are dealing with true monotonic changes in allele frequencies expected of geographic clines shaped by selection within the hybrid zone. When considering all colour loci, *ROS* and *EL* exhibited substantially narrower clines compared to the genomic background (Fig 3; Fig 4). The colour loci *FLA* and *CRE* showed shallower clines than *ROS* and *EL*, but were still distinguishable from the genomic background. These other colour loci were amongst the steepest relative to clines located at similar geographic (centre) positions. For example, filtering only for loci with centres located in the yellow flanks of the hybrid zone between 8-11km, the steepest clines in this region were all clustered around a narrow genomic region on chromosome 5, which we term Chr5CC (Chromosome 5 Cline Cluster). Clines within this region displayed an average width of 7.52km (±1.04km SD), narrower than the background clines in this region 10.24km (±0.91km SD). Similarly, in the core region, the *CRE* locus involved in yellow pigmentation displayed an average cline width of 7.52km (±1.04km SD), substantially narrower than the other clines in this region 10.24km (±0.91km SD) (Fig 4b). The remaining known loci, AUN and SULF showed no clines with the *FastClines* estimates. For the latter, the *SULF* colour locus was not well covered by poolSeq data due to poor sequence coverage across the known deletion in the *A. m. m. striatum* haplotype, although some KASP markers have been developed flanking this deletion and show steep clines similar to *ROS* (Fig 3a; see Bradley et al., 2017).

We next compare cline width and centre estimates at a subset of six loci from *FastClines* (six demes) to traditional descriptive Maximum Likelihood Estimates (MLE) cline fits for much denser geographic sampling (>100 demes) using KASP SNP genotyping. The MLE cline fits used a polymorphic sigmoid model (see methods; 5-parameters; centre, width, p0, p1, *F_ST_*) and cline properties (width and centre) were compared at loci linked to colour genes *ROS*, *EL*, *SULF*, recently identified genes *FLA*, *CRE*, and Chr5CC. Cline width for MLE fits was broadly consistent with the major grouping of cline width estimates from *FastClines* at *ROS*, *EL*, *CRE*, *FLA* and Chr5CC (circle symbols *FastClines* fits vs. square symbols for MLE cline fits in Fig 3; Table S3). The exception was the *SULF* locus which was considerably shallower for the locus with MLE estimates compared to the majority of clines from poolSeq with *FastClines*. However, this was due to the loci genotyped with KASP being excluded from *SULF* region in the poolSeq due to low Δ*p* and coverage. In comparison to cline widths, the range of cline centres from *FastClines* were overestimated compared with MLE cline fits. The latter method of cline fitting showing a much tighter grouping of centres much closer towards the phenotypic centre (Fig 3). However, the step change in cline centre positions either side of the *FLA* gene were consistent between both cline fitting approaches (*FLA1* locus and *FLA2* locus on Fig 3).

### Geographic clines cluster around known flower colour loci

Divergent loci (Δ*p* ≥ 0.9) with steep clines were found on all chromosomes (Fig 5). Cline width ranged from steep to shallow across all chromosomes (Fig 5a), whereas cline centre tended to display different values across chromosomes (Fig 5b). For example, the majority of background loci across all chromosomes and some loci linked to colour loci (*AUN*, *SULF*) exhibited cline centres shifted towards the yellow flank whereas other loci (*ROS*, *CRE*) were centred towards the phenotype transition or slightly towards the magenta flank. To understand how divergent loci with clines were distributed across the genome, we divided each chromosome into 100kb and 10 kb blocks and counted the number of loci with allele frequency differences (Δ*p*) greater than 0.9 between pools 1 (YP4) and 6 (MP11) that also exhibited clines in each block. Only 3.7% of 100 kb blocks contained at least one clinal SNP. Furthermore, considering only 100 kb blocks with at least one cline, the mean percentage of smaller 10kb windows (within each 100kb block) with at least one clinal loci was 19.3%, indicating clines are often localized (Fig 5c). The main exceptions included one region of chromosome 2, with a ∼500kb block of the genome with 90-100% of 10kb windows within the block containing clinal loci. The other exception was on chromosome 6, with all 10kb windows in one 100kb block containing clines (Fig 5c).

**Figure 5.**
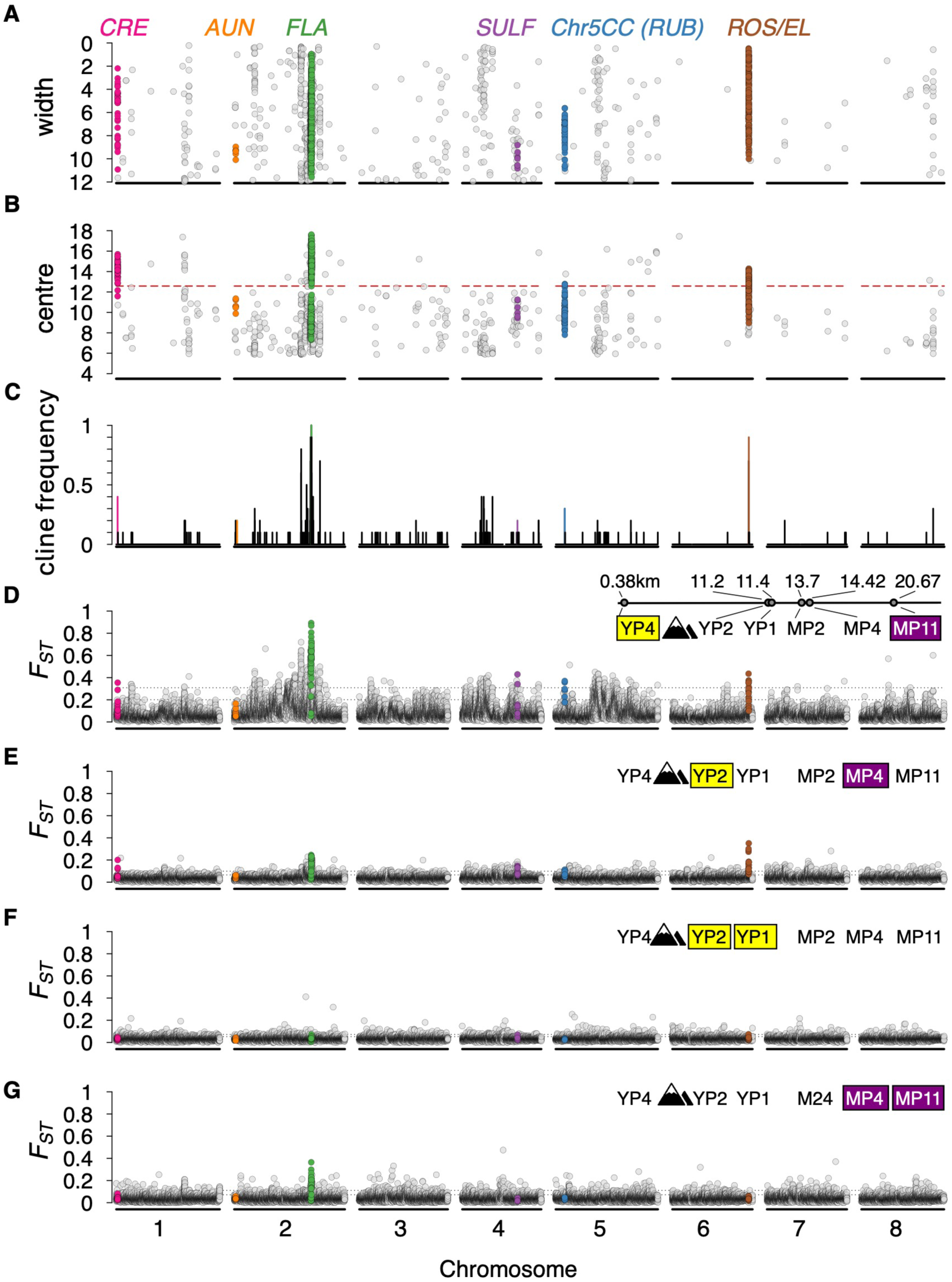
Genome scans between whole genome pools of *A. m. m.* var. *striatum* (yellow) and *A. m. m* var. *pseudomajus* (magenta). Panels show **(A)** cline width with *FastClines*, **(B)** cline centre with *FastClines* **(C)** cline frequency in windows (from *FastClines*) in 10kb windows, **(D)** *F_ST_* pools YP4 x MP11 in 10kb windows (different colours and over mountain pass), **(E)** *F_ST_* pools YP2 x MP4 (different colours and no mountain pass), **(F)** *F_ST_* pools YP2 x YP1 (same colours and no mountain pass), **(G)** *F_ST_* pools MP4 x MP11 (same colours and no mountain pass). Dashed vertical lines indicate phenotype cline centre (red), and 99^th^% and 95^th^% quantiles for *F_ST_* panels (black dashed). Insert in panel (D) indicates relative positions of the six pools along the transect (in km). Window *F_ST_* within 100kb of colour loci are coloured separately for CRE (pink), AUN (orange), FLA (green), SULF (purple), RUB (blue), ROS and EL (brown).

Clinal loci were significantly more clustered in the genome together compared to expectations based on random positions across the genome on chromosomes 2 and 6, but not significantly clustered on the remaining chromosomes (Fig 5a; Table S4). This suggests that some chromosomes exhibit tight concentrations of clines in localized areas of the genome, whereas others exhibit scattered clines along the entire chromosome (Fig 5). Geographic clines, irrespective of properties, were largely clustered in close proximity to the six known genes involved in regulating flower colour. The majority, n = 2918 (76.2%) of divergent loci with clines were located within 300kb of known colour genes (Fig 5). Dense clusters of clines in small genomic regions on each chromosome tagged *ROS*, *EL*, and *SULF*, and recently discovered *CRE* (Richardson et al., in prep), *FLA* (Bradley et al., in prep) and a gene located at Chr5CC (see below). Colour loci were located on the chromosome (Chr) block with the highest frequency of windows with clines on Chr 1 (*CREMOSA*; Richardson et al in prep), Chr 2 (*FLAVIA*; Bradley et al., in prep), and Chr 6 (*ROSEA*/*ELUTA*), except for *SULF* on Chr 4 which was located in the 4^th^ ranked block (Fig 5c).

### Comparison between ***F_ST_*** and cline scans for identifying barrier loci

To understand how pair-wise relative differentiation (*F_ST_*) associates with colour loci and geographic clines, we first examined the number of 10kb *F_ST_* outlier windows (in the 99^th^% quantile) that intersect with colour genes. In contrast to geographic clines, excess *F_ST_* was less consistent in recovering known and recently identified colour loci. Comparing all nine pair-wise populations of yellow and magenta pools, yielded 481 to 694, 10kb windows with excess *F_ST._* Only the *ROS* gene intersected with *F_ST_* outliers in all pair-wise comparisons for at least one 10kb window (Fig 6). The *CRE* gene and *FLA* displayed windows in *F_ST_* outliers in five and six of the pair-wise comparisons, respectively. This was followed by *SULF* present in only two of nine, and both Chr5CC and *AUN* were not present in any pair-wise comparisons. However, for AUN, no windows were in the outlier set (Fig 6). Some windows of excess *F_ST_* were identified even between comparisons of two pools from the same variety (e.g. FLA linked YP4 vs YP2 and MP2 vs. MP11; Fig 6). Permutation tests revealed an enrichment of clinal loci with *F_ST_* outlier windows for most pair-wise comparisons. However, for some pair-wise comparisons many *F_ST_* outlier windows contain no clines. For example, for the most distant populations, YP4 and MP11, only 69% of outlier *F_ST_* windows have clines (Table S4). This percentage of *F_ST_* outlier windows with clines diminished as closer pools of yellow and magenta are compared (e.g. YP3 and MP4 and Δ*p* > 0.9, 27% of *F_ST_* outlier windows has clinal loci; Table S4). Compared to all metrics, *F_ST_* was consistently elevated for windows containing clinal loci compared to windows without clines (Table S5).

**Figure 6.**
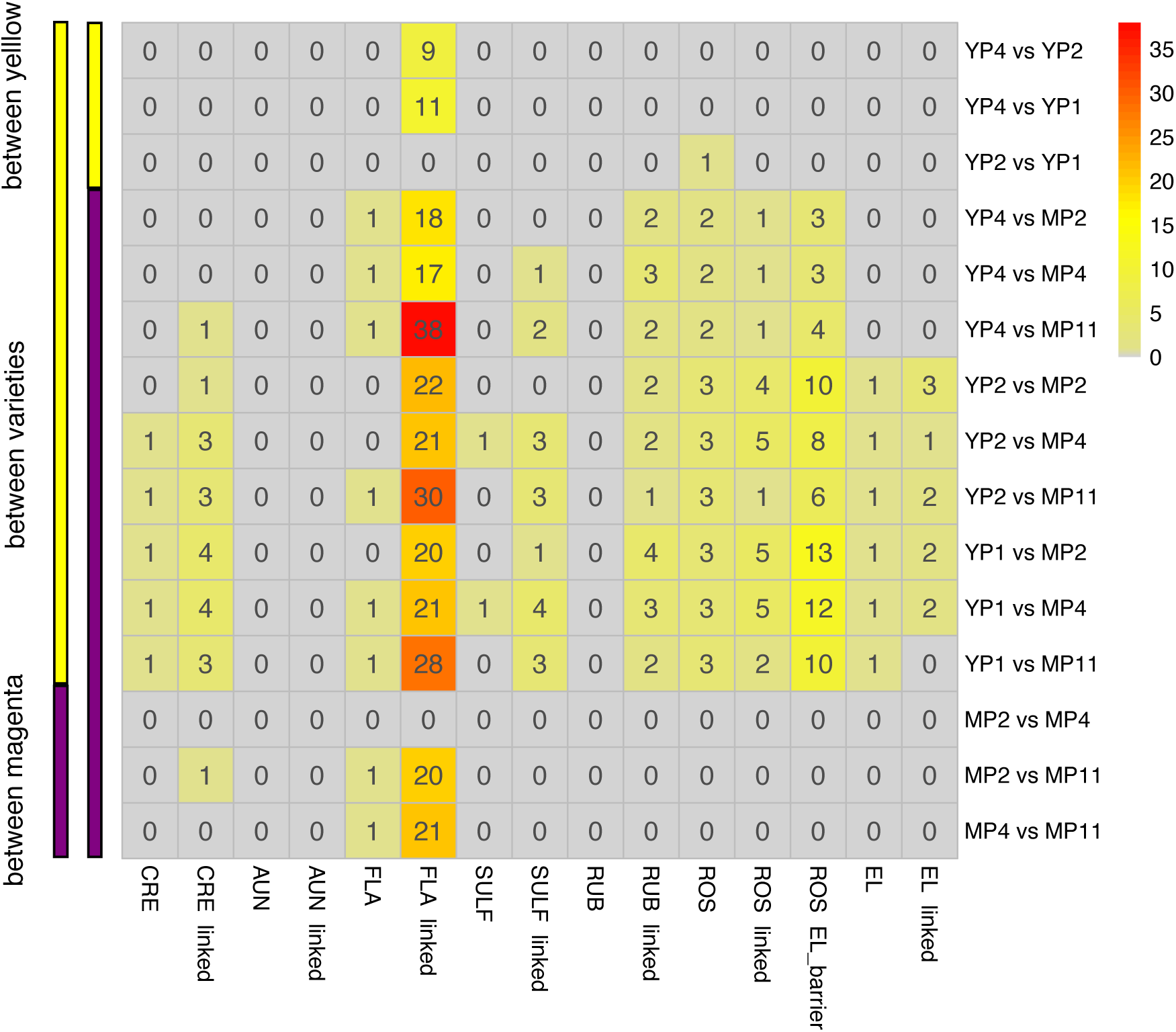
All pair-wise population comparisons of ***F_ST_*** and the number of 10kb windows in colour genes and linked regions along the genome. All pair-wise population pairs shown in rows, with the main colour of the varieties in each pair indicated (left colour bars) highlighting comparisons between yellow and magenta or within variety outliers. Columns indicated the number of windows in the 99^th^ % quantile for *F_ST_* at each of the colour genes and also regions closely linked (<100kb). For the *FLA* gene, closely linked regions are larger (<400kb) due to this being an area of low recombination in the genome. Numbers in each cell indicate how many 10kb windows were detected in the 99^th^ % quantile. Legend displays the colour match for number of outlier windows.

### *RUBIA* locus controls magenta intensity

We interrogated a set of clines from Chr5CC showing the steepest widths in the geographic region left of the core around 9-10km centre position with *FastClines* (Fig 3) and tested for associations between specific genotypes and phenotypes. No genes associated with colour phenotypes had been previously identified in Chr5CC. This region, was the top ranked cluster of clines on chromosome 5, with 87 divergent loci with geographic clines within a tight genomic region spanning 70kb (Fig 5, Fig S9).

To determine whether Chr5CC carried a previously unidentified locus affecting flower colour, we analysed populations derived from crosses between *A. m. m. var. striatum* and *A. m. m var. pseudomajus*. Flower photographs were grouped according to *ROS*, *EL*, and *SULF* genotypes. For each group, we carried out three independent rankings of flowers according to magenta intensity, yielding two bins (Figure 7a). Ranked flowers were also genotyped for SNPs from Chr5CC. Pooling SNP frequencies across genotypic classes showed that the frequency of *A. m. m var. pseudomajus* SNPs for Chr5CC was significantly enriched in the high magenta bin, and depleted in the low magenta bin (*χ^2^* test, *p* = 1.6 x 10^−7^). These results suggest that Chr5CC harbours a magenta flower colour locus, hereafter named *RUBIA* (*RUB*).

**Figure 7.**
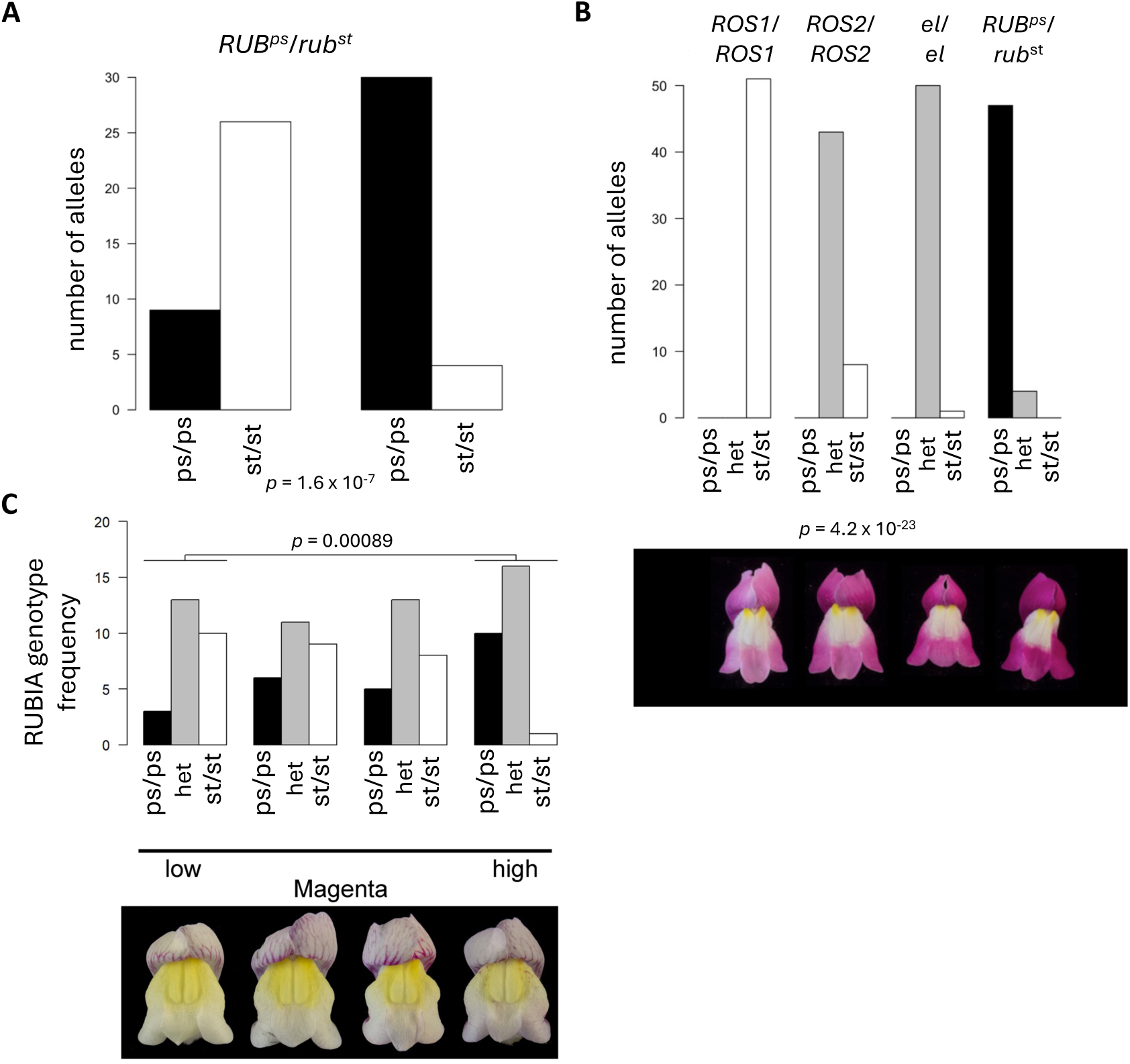
**(A).** Flower colour ranking results for an *F_2_* hybrid population segregating for Chr5CC *F_2_* plants were genotyped for the known flower colour loci *ROS*, *EL*, and *SULF*, and shared genetic backgrounds were grouped. Photographs from different groups were then separately ranked according to magenta intensity, yielding a low magenta bin (left) and a high magenta bin (right). Counts of the *A. m. m. pseudomajus* (ps) and *A. m. m. striatum* (st) Chr5CC homozygotes, summed across all genotypic groups, are shown in black and white. **(B)** Flower colour ranking results for an *F_4_* hybrid population segregating for Chr5CC. Photographs were ranked according to magenta intensity, yielding four quartiles of increasing magenta. Counts of the *A. m. m. pseudomajus* (ps) and *A. m. m. striatum* (st) Chr5CC homozygotes are shown in black and white, with heterozygotes (het) shown in grey. The genetic background of the *F_4_* family is shown above the figure. Representative flower photographs are shown below each quartile. **(C)** Flower colour ranking for an F4 population that was *ros^st^ El^st^/ros^st^ El^st^* and segregating for *RUB* alleles. As in b, the flowers were ranked for magenta and genotypes analysed in quartiles shown with representative flowers of each quartile shown below.

To further test this hypothesis, we genotyped *F_4_* populations that were homozygous at *ROS EL* and *SULF*, but segregating for Chr5CC SNPs and repeated the ranking process. To infer genetic dominance relationships between *A. m. m var. pseudomajus* and *A. m. m var. striatum* SNPs, we split the rank into four quartiles. Ranking according to magenta intensity showed a depletion of *A. m. m. var. pseudomajus* Chr5CC SNPs in the low magenta quartile, and a depletion of *A. m. m. var. striatum* Chr5CC SNPs in the high magenta quartile (*χ^2^* test, *p* = 4.2 x 10^−23^). This suggests that the *A. m. m var. pseudomajus RUB* allele increases magenta (Figure 7b). The two middle quartiles were populated almost exclusively by heterozygotes showing intermediate magenta intensity, suggesting that *RUB^ps^* and *rub^st^* are alleles are semidominant.

To determine whether *RUB* alleles modified pigment intensity in a background carrying striatum alleles at *ROS, EL* and *SULF,* we analysed a population homozygous for *ros^st^ EL^st^* and *sulf^st^* segregating for *RUB* alleles. We ranked the flowers for spread of yellow or magenta and genotyped them for *RUB*. Comparison of the upper and lower quartiles revealed no association with yellow but a significant association of *RUB^p^* allele with high magenta and *rub^st^* with low magenta (*p*=0.00089) (Fig. 7c). Though significant, the effect on magenta was more subtle in this background than in the *pseudomajus ROS^ps^ el^ps^ SULF^ps^* background (compare Fig. 7b with Fig. 7c). Although subtle to the human eye, the effect of a magenta tint in normally yellow regions of the flower may be to reduce colour contrast for pollinators.

This result predicts that *RUB* genotype should correlate with magenta flower colour in plants sampled from the hybrid zone. To test this prediction, we used KASP SNP genotypes at *RUB* together with quantitative colour phenotyping for plants along the transect at the hybrid zone. The allele fixed in *A. m. m* var. *pseudomajus* denoted as *RUB^ps^* and the alternative *A. m. m* var. *striatum* allele denoted as *rub^st^*, although this allele was only at high frequency in yellow population over the pass (Fig 3c). To test for associations at the *RUB* locus, we used linear regressions between colour (HSV space; Hue, Saturation and Value in six floral regions) and the SNP markers linked to *RUB* and *ROS1.* We also included interaction effects between *RUB* and *ROS1* which improved model fits (*F* = 12.9, *p* = 0.00035). The overall model explained a significant amount of variation for Hue (Adj *R*^2^ = 0.49) with a strong effect of *ROS1* (*p* = 1.8 x 10^−09^) but not *RUB* (*p* > 0.05). However, a significant interaction effect between *RUB* and *ROS1* on Hue was detected across central regions of the flower (e.g. Hue 4, *F* = 151.5, *p* = 2.2 x 10^−16^, Fig 8d) (for all floral regions see Table S7). These results suggest an epistatic gene interaction, whereby the *RUB^ps^* allele from *A. m. m* var. *pseudomajus* enhances the intensity of magenta in these floral regions on high magenta backgrounds (*ROS^ps^*/*ROS^ps^* and *ROS^ps^*/*ROS^st^*) but not on low magenta backgrounds (*ros^st^/ros^st^*) (Fig 8d and Fig 8e). However, it is possible that an effect in the low magenta background would have been missed given the subtle signal detected in the crosses (Fig 7c). Also, we cannot rule out the possibility that a recombinant *RUB* allele that is functionally similar to that in *A. m. m.* var *striatum* but that carries *pseudomajus* markers, has introgressed into *A. m. m.* var *striatum,* raising the apparent frequency of *pseudomajus* alleles on the left flank.

**Figure 8.**
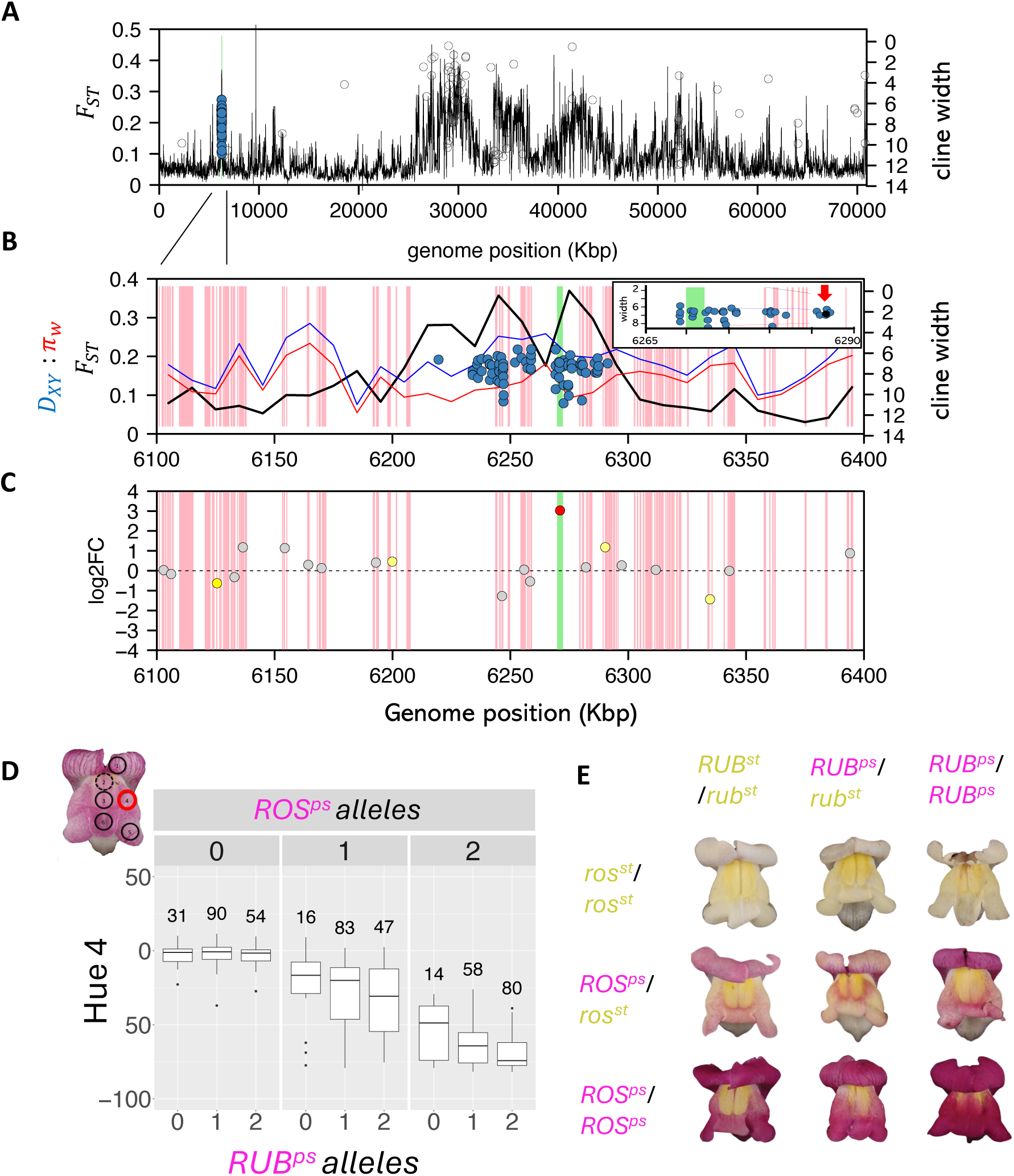
Cline cluster on chromosome 5 in relation to *RUB* locus. **(A)** Genome scan of *F_ST_* (black line), *D_xy_* (blue line) *π*_w_ (red line) along chromosome 5 for pools YP4 vs MP11, **(B)** Genome scan of *F_ST_, D_xy_*, *π*_w_ (YP4 vs MP11) and cline widths (blue circles) from poolSeq for the 300kb region on chromosome 5 (20kb within inset) with the largest cluster of clines on the chromosome (Chr5CC). *D_xy_*, *π*_w_ multiplied by 10. Annotated genes (pink regions), the location of Flavanol Synthase (*FLS*) (green region) and location of KASP SNP marker for *RUB* (red arrow in inset), **(C)** differential gene expression (DGE) from RNAseq for floral tissue of three *A. m. m* var. *pseudomajus* (magenta) vs three *A. m. m* var. *striatum* (yellow) plants indicating transcripts with no significant DGE (grey), weakly significant (yellow) and strongly significant (red). **(D)** colour phenotype scores for Hue in location 4 on flowers for the three *RUB* genotypes (0,1,2) at each of the three *ROS* genotypes (panels, 0,1,2) where numbers represent three genotypes and copy number of *A. m. m* var. *pseudomajus* alleles, **(E)** example of phenotypes with median Hue 4 values for haplotypes at *ROS* and *RUB* from hybrid zone.

At the centre of the chromosome 5 region carrying *RUBIA* was a gene encoding flavanol synthase (*FLS*) a key enzyme involved in the Anthocyanin colour pathway (Fig 8b). The SNP marker used for MLE clines and in genotype-phenotype associations was ∼150kbp upstream of *FLS* with eight annotated genes identified within this interval (Fig 8b inset). The *FLS* region also exhibited slightly elevated *F_ST_*, with some of the linked windows showing higher *F_ST_* (depending on which yellow and magenta pool compared; Fig 6). Elevated *F_ST_ was* not strongly driven by low diversity (*π*) or excess divergence (*D_xy_*) (Fig 8b).

Next, we aimed to confirm whether any genes in this region displayed differential gene expression linked to phenotypic differences. Using RNAseq data from corolla (petal) tissues from two sets of yellow and magenta plants sampled from seed in the flanks of the hybrid zone, transcripts were compared for differential gene expression across the genome. Sorting of the transcripts by the degree of differential expression revealed that the sequences in tight linkage with the *FLS* gene displayed the highest differential gene expression levels on the entire chromosome (Fig 8c). This finding is consistent with *FLS* being responsible for tissue specific differences between the subspecies, in this case colour pigments as the only trait that clearly differs between the two groups. Here, expression of *FLS* is high in *A. m. m* var. *striatum* (yellow) and low in *A. m. m* var. *pseudomajus*.

## Discussion

### An efficient alternative for surveying geographic clines across genomes

Our new approach for estimating genome wide geographic clines has enabled a rapid genome wide perspective on cline properties. In comparison to traditional Maximum Likelihood Estimate (MLE) cline fitting, the *FastClines* method has a significant advantage in that its very fast and doesn’t require numerical optimization using simulated annealing procedure (e.g. Derryberry et al., 2014). Another advantage of the method is the ability to obtain reasonable estimates of cline properties with as few as six demes along a transect through a hybrid zone. Although this method provides an efficient alternative to traditional cline fitting, several caveats can be distinguished. First, for computational speed, this method defines cline centre by median mass of allele frequency and cline width via total heterozygosity. However, these estimates could be different to the steepest point and the inverse of the steepest gradient from a sigmoid geographic cline. These differences may be especially pronounced when geographic clines become significantly asymmetric. Secondly, although estimates can be calculated with few unevenly spaced demes, this will likely introduce biases in the positions of cline centres and widths. However, some of these challenges can be overcome by simply increasing the number of demes and validation by estimating cline properties using traditional MLE fitting at some representative loci.

For the *Antirrhinum* hybrid zone, our comparison of cline fitting methods indicated that *FastClines* produced a narrowing of clines widths when cline position (centre) moves towards the edges of the transect. This is likely explained by the coarse spatial sampling (i.e. six pools), where width and centre can be confounded, resulting in narrowing of clines due to step changes in frequency between the outer two demes. Simulations of *FastClines* estimates generated this same artefact, but only with small numbers of pools (i.e. ≤8 pools). The accuracy of *FastClines* increases with spatial resolution with greater number of demes along a transect, a feature of empirical studies which will become easier as sequencing costs for whole genome sequencing large numbers of individuals becomes more cost effective. For *Antirrhinum*, inspection of individual loci frequencies also show many of these loci exhibit sharp allele frequency step changes between pool YP4 and YP2 rather than a sigmoid-like change over geographic space. Between these pools is a high mountain pass with no *Antirrhinum* (Fig 1), together with the significantly different *F_ST_* landscape when comparing YP4 with all other pools in Planoles valley (Tavares et al., 2018) suggests the far outer populations on yellow side has little genetic connectivity with the rest of the hybrid zone and this mountain pass represents a geographic barrier to gene flow.

### Clines detect known barriers along the genome

We demonstrate how a genome wide perspective on geographic clines can be a useful approach for narrowing down areas that contain barriers. In the hybrid zone, cline widths of loci linked to most of the colour genes were also narrower than the genomic background (i.e. >100kb away from known genes). This is consistent with the premise that these variants of *Antirrhinum majus* are primarily distinguished by small numbers of large effect genes. The majority of clines were clustered in tight physical linkage near these major effect genes controlling flower colour. The overall low frequency of fixed differences and tight clusters of clines in narrow genomic regions is also consistent with ongoing gene flow and homogenization across most of the genomes (Tavares et al., 2018) with only a small number of barrier loci remaining intact in the face of gene flow. The pattern at many background loci (unlinked to colour loci) exhibiting a sharp allele frequency step over the mountain pass (chromosomes 3, 7, 8), may be a signal of an introgressive sweep of magenta alleles into yellow flanking populations that has been blocked at the mountain pass. This supports theory that predicts that divergently selected alleles are expected to resist introgression and maintain steep clines, whereas neutral or advantageous alleles will exchange freely across the hybrid zone and eventually flatten out. Theory also predicts that equilibrium should be reached quickly for selected loci (*t*=1/*s*) (Barton & Gale, 1993; Barton & Hewitt, 1985). Considering the age of the hybrid zone is at least 100 generations old (Tavares et al., 2018), even the most weakly selected colour loci should have reached equilibrium.

The total number of clines and how they are distributed along genomes remains a challenging property to survey even for whole genomes. For the *Antirrhinum* hybrid zone, we intentionally placed three whole genome pools to the left and three to the right of the flower colour transition. This approach likely contributed to *FastClines* estimates retrieving each of the major flower colour loci and identification of a new locus. Yet we still have very different power to detect clines if they are centred close to the floral phenotype transition, compared to clines that are displaced geographically elsewhere. The second major challenge is how to interpret differences in cline density along the genome. For example, we found large differences in the number of clinal loci around the *FLA* locus compared to the next largest cluster of clines around the *ROS* locus. Given local recombination rates have been estimated as significantly lower at *FLA* compared to *ROS* (Bradley et al in prep), this suggests the local recombination landscape in which selected loci are embedded is important for interpreting cline density. For a barrier locus, lower recombination rates increase the extent of linked selection on neutral loci, slowing the exchange of alleles through recombination between neutral and barrier loci across the genome. However, even with comparable recombination rates, the number of individual tightly linked barrier loci will influence how many clines are detected. Theory predicts the effects of linkage around a single site to be more localised than for a region that contains two or more tightly linked barrier loci, as multiple loci generate a stronger barrier (Tavares et al., 2018) and hence influence the number of clinal loci surrounding the causal loci. Therefore, expectations on the numbers of clines along the genome is complex and by itself not a good indication of the importance of the area as a barrier. Interpreting cline density will require careful consideration initial divergence (Δ*p*) local recombination landscape and the genetic architecture of barriers.

Differences in the geographic position of clines at independent barrier loci also provide important insight into the role of gene interactions. With both anthocyanin and aurone pigmentation interacting to constitute the parental phenotypes (magenta vs yellow), theory predicts these loci will be synergistically coupled (concordant cline centres) upon secondary contact due to the influx of parental gene combinations and linkage disequilibrium (Barton & Hewitt, 1985). Although there is some evidence of different cline centres, the range of cline centres were likely overestimated with *FastClines* due to low numbers of demes the coarser nature of estimates from this approach. Spatially dense sampling and testing of alternative cline models indicate that the clines at these colour loci are mostly coincident yet display stepped clines with different asymmetrical tails (Surendranadh et al in prep). Assuming a symmetric sigmoid cline model when asymmetries either side of clines exist, are known to incorrectly push cline centre estimates apart (Baird & Macholan, 2012). This is one limitation of the *FastClines* method as it assumes a polymorphic yet symmetric sigmoid cline, suggesting some of the spread of cline centres observed may be driven by these asymmetries. However, one exception is at the loci either side of *FLA*, a recently described gene that interacts with *SULF* to determine yellow pigmentation (Bradley et al. in prep). Closer inspection of cline centres walking along the chromosome near the *FLA* gene revealed a step change in cline centre position directly over the gene region. Traditional MLE cline fitting confirms a different cline centre estimate at representative loci either side of this gene (Surendranadh et al in prep). This step change in centre estimates coincides with the breakpoint of a recombinant haplotype at the *FLA* locus which has a different phenotype (Bradley et al in prep). Despite some of the caveats of this approach, these results highlight the value of *FastClines* and the potential for detecting course step changes in cline properties to flag potentially important genomic areas related to phenotypic differences.

### Clines narrow down the RUBIA locus that modifies magenta intensity

We demonstrate that focusing on clusters of steep clines along the genome can be useful for identifying new barrier loci contributing to phenotypic trait divergence. This geographic cline approach helped locate a new gene *RUB* which *F_ST_* and other diversity metrics did not as readily detect. This *RUB* locus is tightly linked to *FLS*, with RNAseq showing expression of *FLS* is high in *A. m. m* var. *striatum* (yellow) and low in *A. m. m* var. *pseudomajus*. One possible explanation is the *A. m. m. var. striatum* variants at FLS diverts common substrates into flavanols, so reducing the flux into anthocyanins (Aida et al., 2000; Luo et al., 2016). Although the overall level of differential gene expression is not as strong as seen at other loci such as *FLA* (Bradley et al in prep), this may reflect that this region controls a phenotypic difference of smaller effect on pigmentation.

Cline fitting of a polymorphic sigmoid cline using a MLE simulated annealing approach revealed a steep cline fixed (best model, *p* = 0.96) on the *A. m. m* var. *pseudomajus* (magenta) side of the hybrid zone yet the locus remains still polymorphic on the yellow side of the hybrid zone (*p* ∼0.4) (Fig 3). Although the allele is close to asymptote on the far yellow side, this part of the transect is in another valley and not well connected to the rest of the hybrid zone (Tavares et al., 2018). Therefore, the *RUB* locus on the yellow side of the hybrid zone exposed to gene flow is largely polymorphic in contrast to the magenta side (i.e. excluding areas over the mountain pass). This could be due to the *RUB* allele from *A. m. m* var. *pseudomajus* having limited phenotypic effect on predominantly *A. m. m* var. *striatum* (yellow) parental genotypes and thus is unseen by selection in areas where yellow backgrounds dominate. Future efforts to test the strength of selection of alternative alleles on either side of the hybrid zone will be important in revealing if the epistatic interaction between *RUB* on different *ROS* backgrounds translates to fitness epistasis.

### Conclusions

Our study highlights several future challenges for research in understanding genome-wide distributions of clines. Firstly, the covariance of cline parameters due to tight linkage in *Antirrhinum* highlights the need for future theoretical approaches that deal with the issue of non-independence among loci. Unlike classical hybrid zone studies using small numbers of markers, the move to whole genomes necessarily results in loci in tighter linkage and thus many of them will not be statistically independent. This is also an issue for other genomic cline methods, which are based on detecting outliers from the neutral background loci, which are wrongly presumed independent. One approach to dealing with this problem is to move beyond single loci and follow haplotype blocks through the hybrid zone (Sedghifar et al., 2015). Secondly, we still lack a theoretical understanding of the expected distribution of clines across genome wide data and how the rate of false positives depends on the time since secondary contact, drift, local recombination rate and the strength of selection. The much higher frequency of clinal loci around the *FLA* gene (region of low recombination), compared with *ROS* and *EL* (high recombination), highlights the importance of the recombination landscape and why the density of clines in a region is not a good indication of the importance of the region as a barrier. This issue is similar to the elevated *F_ST_* associated with regions of the genome with lower rates of recombination and nucleotide diversity (Cruickshank & Hahn, 2014b; McGaugh et al., 2012). Regardless of the evolutionary forces responsible for the signal, more clines may be expected in areas with low diversity and lower rates of recombination and these features need to be considered when interpreting the density of clines along chromosomes. Determining how far genetic barriers persist at neutral sites around a selected locus that factors in these genomic features will also provide insight into the processes structuring genome wide divergence. This will enable the further dissection of the role of stochastic, historical and contemporary forces in driving patterns of divergence and clines across genomes.

## Methods and Materials

### *Antirrhinum* hybrid zone sampling

In order to conduct genome scans and estimate geographic clines with *FastClines*, we subsampled individuals from six demes along a 1-dimensional transect at the hybrid zone near Planoles, Spain (Fig 2). The six demes were arrayed along a 1-D transect spanning ∼24 km (following Bradley et al., 2017), with tighter spacing of demes near the centre of the phenotypic cline to capture the steep transition. To ensure spatial coverage of the flower colour cline, three subpopulations were selected in the predominately yellow regions west of the centre of the cline and three in magenta dominated regions to the east (see Fig 1 and Table S1 for locations). In the outermost populations (YP4 and MP11) only yellow and only magenta individuals are present, respectively. However, in the remaining (YP2, YP1, MP1, MP4) hybrids and both parental phenotypes are present (Fig 1). Altitude gradually increases up the valley going West, above ∼1600 metres in altitude, *Antirrhinum* is absent. Thus, a break of ∼3 kilometres in the distribution of *Antirrhinum* plants coincides with a mountain pass. The outermost yellow population (YP4) is situated west of this pass, while the other five population samples are east of the mountain pass (Fig 1).

Each of the six demes included a random sub-set of n = 50 individuals within a 50 metre radius from a larger sample. In this larger project, over 30,000 plants were located to within 3.4 metres with a GPS (Trimble GeoXT datalogger), leaf tissue collected for DNA extraction and one flower taken for phenotyping (see details in Ringbauer et al., 2018). Following Whibley et al., (2006), individuals were categorized into six phenotype/genotype classes on the basis of anthocyanin and aurone pigmentation across the flowers.

### Whole genome sequencing and KASP SNP genotying

We sequenced the six pools and used whole genome sequencing (n= 50 individuals in each pool) carried out using Illumina HiSeq. The DNA extraction methods, sequencing and bioinformatic pipelines and filtering are described in detail previously (Tavares et al., 2018), with the exception that the original data has been re-mapped to *Antirrhinum* reference genome v3.5 (Li et al., 2019; Zhu et al., 2023).

The larger sample of individuals (n = 30,000) were also SNP genotyped with Kompetitive Allele Specific PCR (KASP) (following Ringbauer et al., 2018) as part of a separate pedigree project. Here, we used this KASP SNP genotype dataset for genotype-phenotype associations (see below) and to fit descriptive clines and verify FastCline estimates from poolSeq. Candidate loci for the KASP marker design were identified from the poolSeq data including loci for (i) each gene known to influence flower colour (*ROS*, *EL*, *SULF*, *FLA, CRE*), (ii) major cline clusters identified (using *FastClines*) on each chromosome (including new colour loci *RUB*)(see Table S2 for marker details). All genotyping and scoring was carried out by LGC genomics.

### Genome wide estimation of geographic clines properties using FastClines

Estimating geographic cline parameters for whole-genome data is not straightforward, especially when alleles are not fixed for alternative alleles in the most outer populations (Δ*p*_1,6_ ≠ 1.0) when many parameters need to be fitted for thousands of loci across whole genomes. One approach is the cline approximation method described by Polechova & Barton (2011), which is not computationally demanding and provides a reasonable approximation to detailed cline model fitting (Gay et al., 2008; e.g. Szymura & Barton, 1986). They showed that the width of a cline can be approximated for diagnostic loci as the integral of heterozygote frequencies over space. For discrete demes, this is essentially twice the sum in the frequency of heterozygotes across all demes,

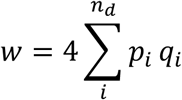

where, *n_d_* is the number of demes, *p_i_* and *q_i_* the allele frequencies in the *i*th deme, and width is measured in deme spacing. Similarly, the centre of the cline can be estimated as the sum of allele frequencies adjusted by half a deme,

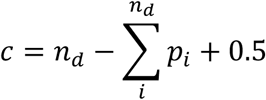

However, with real data loci may not be fixed for alternative alleles in one or both parental populations. Moreover, this model assumes equal spacing among demes.

We developed a new method we call *FastClines*, which extends this approach to non-diagnostic loci that also accounts for differences in deme spacing (size) across a hybrid zone (see Figure 1 for example). For diagnostic or non-diagnostic loci, we denote the allele frequencies in the two parental populations (or outer most demes) as *p*_0_ and *p*_1_ and assume that they are known. We then restrict the integral to lie within the interval of the allele frequency differences between the parental populations for each deme as (*p_i_* – *p*_0_)(*p*_1_ − *p_i_*). The cline width can then be approximated by,

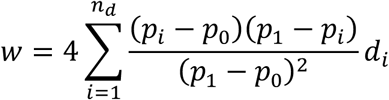

where *d_i_* is the span of the deme, and equal to half the Euclidean distance between the midpoint of the samples in each deme. For the outermost demes we extend this outwards to the same distance. This term *d_i_* provides the appropriate scaling to account for irregular spacing of demes (see Supporting information S1 for example).

Similarly, cline centre is approximated by,

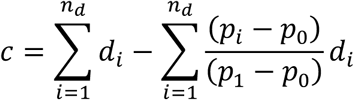

where the first term orientates the centre with respect to *p*_0_.

We use this novel *FastClines* method on the *Antirrhinum* data, taking the pooled allele frequencies as given across the hybrid zone. Here the demes were irregularly spaced apart and we scaled according to the midpoint distance between the demes mid point [*d_i_* = (6000,5000,1500,1400,3500,6000); Table S1] along a transect which was set from the descriptive cline fitting (see below). We include only loci with strong allele frequency differences between the outer pools Δ*p*≥ 0.90 and filter out rare negative cline widths (Text S1). All estimates were calculated in a custom Python script *FastClines* for cline approximations for whole genome data (https://github.com/dfield007/fastClines).

### Descriptive cline fitting with simulated annealing

We fitted traditional descriptive clines using SNP genotypes to compare with FastClines estimates from poolSeq. We settled on 200m demes, this scale minimized deficits from Hardy-Weinberg equilibrium whilst ensuring sufficient sample size within demes (mean = 40, sd = 20, range 10-200) and yielded ∼35,000 individuals across ∼400 demes across the hybrid zone (i.e. numbers vary slightly across loci). Only one marker within *ROS* displayed significant departures from HW with a deficit of heterozygotes (*F*>0) even at small spatial scales (<50 metres). Therefore, we expect based on theory that this feature may generate narrower cline widths than expected. A more detailed examination of the effects of deme size and departures from HW is presented elsewhere (Surendranadh et al., in prep).

The clines were characterised for each colour locus, using a modified version of a custom R script (https://github.com/dfield007/slowClines) described in Bradley et al., (2017) and Tavares et al., (2018). This script uses fits a symmetric polymorphic sigmoid cline with five parameters:

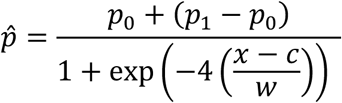

Where *c* = cline centre, *w* = cline width (1/gradient), *p*_0_ = allele frequency at the asymptote in the west (*A. m. m. var striatum* parental allele frequency) and *p*_1_= allele frequency at the asymptote in the east (*A.m. m. var pseudomajus* parental allele frequency) and 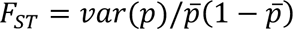. For the *F_st_* parameter, we fitted a beta-binomial error term to account for the variance in allele frequencies across demes and to control for population structure along the cline. Prior to commencing cline fitting algorithm, the larger data set was randomly thinned within each deme to reduce the computational time, resulting in 10,000 individual genotypes across ∼120 demes.

We use a metropolis-hastings (simulated annealing) algorithm to sample the likelihood surface of the cline fit. We begin the algorithm with a random set of parameters which are changed randomly and the log likelihood *logL* is computed at each iteration. When the next iteration *logL’’* has a greater log likelihood than the previous likelihood *logL’* (i.e. *logL’’* > *logL’*), the new parameters are accepted. If the next iteration is lower (*logL’’* < *logL’*), we accept with a probability *logL’’*/*logL’*. To ensure ample exploration of the likelihood surface, the jump size for the next set of parameters are adjusted by a factor of 1.05 when accepted (accept scale) and by (1/1.05) when parameters are rejected (reject scale). After some tests of different accept and rejection scales, we found these values achieved efficient mixing and exploration of the likelihood surface with an acceptance rate ∼0.5. This algorithm was run for 50,000 iterations with a burn-in = 2000. From this we find the most likely cline parameters, maximum log*L* and assume the likelihood surface follows a chi-square distribution to find −2 log*L* max 95% credible regions. We visually inspected the joint likelihood surface for each run. Each run was repeated with randomly chosen starting parameters to ensure reproducibility. Different cline transect orientations were tested (Fig S3 and Fig S4). More detailed cline fitting of alternative cline shapes and asymmetries combined with estimates of selection are being address elsewhere (Surendranadh et al., in prep).

### Simulations of cline estimates from FastClines

We used simulations to examine the power of the *FastClines* approximation method to distinguish coarse shifts in the centre and width of clines. Rather than forward simulating the dynamics of hybrid zones through generations following dispersal and selection acting on loci (e.g. Westram et al 2018), this simulation assumes smooth clines with known properties are already in place at a particular fixed time point. The position and width of clines is expected to vary from the true parameters due to the demographic process (e.g. drift that affects allele frequencies, non-random mating) and sampling effects (number of demes, sampling error, variation in sequencing depth and sequencing error). We modeled each of these factors using a similar setup to the whole genome data for *Antirrhinum* and explored other arrangements with more sampled demes.

The setup follows a 1-dimensional landscape with a symmetric polymorphic sigmoid cline with five parameters (see cline model above). For simplicity, the total transect length matched the *Antirrhinum* hybrid zone (deme positions from 0 to 24km). The simulation begins by generating a matrix of allele frequencies in demes along columns (demes along the cline transect) with replicate sampling along rows. For each deme, observed allele frequencies are drawn from a beta distribution where alpha = (1/ *F*_ST_)-1). In each deme, the expected allele frequency is based on the cline parameters yet the observed values fluctuate, first due to drift following a Beta distribution,

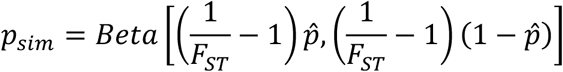

where *F*_ST_ = 0.03 (following estimates at neutral loci for *Antirrhinum*). Next, a specified number of individuals to represent sampling from field sites as is each deme were drawn for each of the possible homozygote diploid genotypes (*g_kk_*, *g_jj_*) and heterozygote (*g_kj_*). The diploid genotypes were randomly drawn from the observed allele frequencies 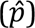 following a binomial and departures from random mating expectations at Hardy-Weinberg equilibrium (HW) modeled by varying the *F* parameter below for each of three genotypes as,

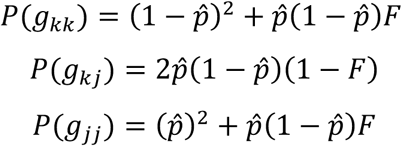

To simulate variation in sequencing depth, we used a normal distribution with a mean and variance calculated from the poolSeq data. The method draws sequencing reads (with replacement) from the alleles contained in the diploid genotypes in each deme. We used a normal distribution with a mean and variance following the observed values in the *Antirrhinum* aligned whole genome data with sequencing errors introduced at 1 x 10^−4^. For each set of parameter values we ran replicate simulations (*n*= 10,000) to examine the variation in cline width and centre estimate.

We first explored parameter space with a combination of width and number of demes to test how cline width is improved with density of geographic sampling [w={2,4,6,8}, demes={6, 8, 10, 12}]. We also tested varying both width and centre values [w={2,4,6,8}, centre = {8, 10, 12, 14, 16}]. To examine the effect of departures from HW and variance in allele frequencies due to structure, we ran simulations for one set of cline parameters (c = 450, w = 150) for a range of *F* = {0,0.5,0.1,0.2} and *F*_ST_ = {0.01, 0.05, 0.1}. Similarly, to examine the effect of increased sequencing depth, we also ran this set of cline parameters for a range of depth (25, 50, 100) with variance similar to observed values (*σ*^2^ = 5). Scripts for cline simulations were written in R and are available online (https://github.com/dfield007/fastClines).

### Genomic diversity and divergence

We quantified relative genetic divergence and diversity within and between the six pooled populations within 10kb sliding windows using the methods outlined in Tavares et al., (2018) with a few modifications. These included remapping the data to the *Antirrhinum* reference genome v3.5 and running through the custom SlidingWindows script v1.14 (https://github.com/dfield007/slidingWindows). We estimated diversity within (*π _w_*) each of the populations, and total diversity (*π_t_*), relative (*F*_ST_) and absolute differentiation (*D_xy_*) between each pair of populations using approaches outlined in Tavares et al., (2018).

### Annotation of genes corresponding to gene clusters

To determine whether the identified clines correspond to phenotypic effects, we annotated protein-coding genes from major cline clusters on each chromosome (top-ranked cline proportions; Fig. 5c). We first extracted cluster-associated genes from the genome annotation file (reference genome version 3.5). We performed a functional annotation using the eggNOG-mapper v2, combining orthology assignments, functional descriptions, and gene ontology (GO) terms (Huerta-Cepas et al., 2019). Simultaneously, the genes were subjected to similarity searches using the BLAST online tool (Basic Local Alignment Search Tool) against the NCBI non-redundant (nr) database. The functional information and gene hits obtained from both eggNOG-mapper and BLAST were manually curated to ensure accuracy and relevance. The location of all known genes known to influence flower colour was combined with genes identified around clinal loci. Genes were then categorized into functional groups, including the broad term ‘colour related gene’ to include all of those involved in the flavonol biosynthetic pathway or known to regulate the expression (intensity or distribution) of colour pigments across parts of the flower in *Antirrhinum majus*.

### Clustering in the genome

We used permutation tests to determine if clinal loci were more clustered in the genome than expected by chance. We randomized the labels associated with all SNP positions (i.e., clinal SNP = Y or N) in the genome and calculated the mean distance in bp between all possible pairs of clinal labels. This was repeated 9,999 times, and each time the permuted mean pairwise distance was retained to generate a null distribution. The observed value of the main pairwise distance was calculated for the observed data. We calculated a p-value for the observed value as the number of permuted datasets where the mean pairwise distance was equal to or greater than the observed estimate + 1/number of permutations +1.

### Genotype-phenotype associations at cline cluster chromosome 5

To examine the effect of genomic region on Chr 5 (Chr5CC = RUBIA) which gene annotation discovered was in close proximity to Flavanol Synthase (FLS) we examine genotype-phenotype associations in controlled crosses and wild plants from the hybrid zone. First, we generated an F2 cross between purebred *A. m. m var. pseudomajus* (magenta flowers) and *A. m. m var striatum* (yellow). A sample of n = individuals were SNP genotyped using KASP to confirm genotypes at known colour loci and the clinal SNPs on Chr 5.

In addition, we phenotyped and genotyped plants from the hybrid zone to examine the effect of clinal loci on Chr 5 on flower colour. In the hybrid zone, we randomly selected a subset of the six major colour phenotypes (Magenta, Yellow, Pink, Weak Orange, Full Orange and White; following Whibley et al., (2006)) that has been genotyped with KASP. Using photographs of the front of the flower color measurements were taken in ImageJ (http://imagej.nih.gov/ij/) at six standardized locations on the flower (Fig S14). For each region of the flower, we calculated the mean and standard deviation of Hue, Saturation (Sat), Intensity (Int) and Greyscale (GS) and transformed these values to HSV colour space. The effects of *ROS1* and *RUB* genotypes on colour phenotypes were assessed with linear regressions, with interaction effects included to test for epistasis (see full details Text S4).

### Differential gene expression

RNA was extracted from corolla tissue from snapdragon flowers for three biological replicates of a representative *A. m. m* var. *pseudomajus* (magenta flowers) and *A. m. m* var. *striatum* (yellow). We used the annotated *Antirrhinum* reference (Accession number GWHBJVT00000000) genome v3.5 (http://bioinfo.sibs.ac.cn/Am/) and added “decoy-aware” index for the mapping. For each of the samples we next used Salmon to align and quantify transcript abundance to our annotated genome, including flags to increase the stringency of the mappings (--validateMappings) and to learn and apply corrections for GC bias and primer bias (--gcBias --seqBias). We then performed differential gene expression analyses using Wald test hypothesis in DESeq2 (Love et al., 2014)(implemented in Bioconductor v3.2 in R) with the lfcShrink function to compensate for inflated log2fold changes in genes that have low counts (after filtering out genes with < 10 reads). As a reference point, other known colour genes in the top clusters on Chr 6 (*ROS*/*EL*) and Chr 2 (*FLA*) we found to have the highest outliers for differential gene expression

## Acknowledgements

We are grateful to Annabel Whibley, Monique Burrus, Christophe Andalo, Tom Ellis, Parvathy Surendranadh and members of the Barton group for interesting discussion. We are also grateful for numerous undergraduate volunteers who assisted with collecting of flowers and leaf samples in the field.

