## Supplementary material for "Genome-wide cline analysis identifies new locus contributing to a barrier to gene flow across an *Antirrhinum* hybrid zone"

This SM material contains:
Supplementary Text S1 to Text S5
Supplementary Figures Fig S1 to Fig S18
Supplementary Tables Table S1 to Table S8
References

### Supplementary Text

#### S1: FastClines setup

*Deme locations along transect*

We used the whole genome poolSeq data and the transect positioning through the hybrid zone as described in Tavares et al., 2018. The geographic positions of each pool were located along this transect and collapsed to one dimensional geographic distance (in kilometers) along the transect (Fig 1). The total geographic distance along the transect was set by the larger SNP KASP genotype data with demes located from 0 km to 25.176km. For the poolSeq, the deme location was set as the average Easting and Northing of individuals in each pool (Table S1) with positions then collapsed along the 1-dimensional transect to the following positions [YP4 pool 1: 0.382km, YP1 pool2: 11.227, YP2 pool3: 11.404, MP4 pool4: 13.767, MP4 pool5: 14.424, MP1 pool6: 20.670km].

*Deme spans*

In the fastClines method, we need to define the geographic span of each deme ($d_i$ and see Fig 2). If the demes are equally spaced across a transect, one approach is to use half the distance either side of the geographic centre of each deme as the span of each deme. However, the patchiness of individual samples in natural populations, including the snapdragon system, often preclude setting equal distances between pools. Therefore, to account for possible influence of deme span selection on cline properties, we used the midpoint edges: where the outer edges (pool 1 and 6) were defined by the dimensions of the KASP sample demes (0 to 25.176km) and all internal edges set as the midpoint distances between each pool along the transect [deme edges: 0, 5.423, 11.316, 12.586, 14.095, 17.548, 25.172km] from which the geographic span of each deme was calculated [pool 1: 5.805km, pool2: 5.893, pool3: 1.270, pool4: 1.500, pool5: 3.453, pool6: 7.251km]. The total span of the demes was 25.172km and the midpoint between pool 3 and 4 (transition across centre of phenotype cline and ROS1 gene) was fixed at 12.586km. This allowed for comparisons of cline centres and widths relative to the known position of the ROSEA locus (~12.5km – 13.5km) and the main phenotypic cline.

#### Filtering and cline estimates for *Antirrhinum*

We use this method on the *Antirrhinum* data, taking the pooled allele frequencies as given across the hybrid zone. Here the demes were irregularly spaced apart and we scaled according to the midpoint distance between the demes [$d_i$ = (5800,5890,1200,1500,3450,7200)]. Although allele frequencies estimate may benefit from incorporating errors for pooled data (e.g. Lynch *et al.* 2014), we found a high correlation between allele frequencies estimated from the pools and those estimated from individual genotyping of the same 50 individuals in each pool (see Tavares et al 2018). We include only loci with strong allele frequency differences between the outer pools $\Delta p_{1,6} \geq 0.80$ and $\Delta p_{1,6} \geq 0.90$. For these loci, allele frequencies were polarized so that they increased from West to East (yellow to magenta, respectively). One limitation of this method is that cline reversals (e.g. $p_5 > p_6$) for non-diagnostic loci can result in negative cline widths (Supporting Info xx). However, we only identified three such loci with significant reversals. Considering this occurred at loci with significantly lower depth in the parental populations than the average, this may be due to sampling artifacts and therefore we removed these from further analyses.

Initial analysis using FastClines found n = 12,936 clinal loci with a low depth filtering threshold and allele frequency difference between the outer most pools (min depth x = 10 in at least 5 of 6 pools, $\Delta p_{1,6} \geq 0.80$). This was reduced to n = 10,912 and n = 7,271 loci when increasing filtering to a minimum of 15 and 20 depth, respectively (in at least 5 of 6 pools)

With this method, negative cline widths were detected at n = 299 loci of 12,936 (2.3%), n = 249 loci of 10,912 (2.3%) and n = 162 loci of 7,271 (2.2%) loci when filtering for minimum sequencing depths of 10, 15 and 20, respectively [Figure S13(a -e)]. The majority of loci with negative cline widths were positioned with centres at the extreme ends of the transect. For example, considering loci with at least 15 depth (Figure S13c), we found n = 232 (93.2%) were centred to the extreme left (<6000m) or right (>15,000m) of the transect.

Filtering to loci with $\Delta p_{1,6} \geq 0.90$ had a stronger impact than filtering for higher depth in reducing the frequency of negative cline widths. For example, at minimum depth 15, increasing to $\Delta p_{1,6} >$ 0.9 resulted in 16 loci of 3,826 (0.04%) with negative cline widths compared with 249 loci of 10,912 (2.3%) of loci for $\Delta p_{1,6} > 0.8$.

Negative cline widths were observed for loci with reversals in allele frequencies along the transect. For example, when considering loci with $\Delta p_{1,6} \geq 0.80$, the n = 16 negative cline widths positioned near the hybrid zone centre (10,000 – 15,000m) tended to display allele frequency reversals in the outer first pool (Figure S14).

In summary, these negative clines likely represent artefacts due to the limited number of pools at lower allele frequency differences. This tends to draw widths downward as centres move to the edge of the last deme. Focusing on loci with fixed differences or greater allele frequency differences ($\Delta p > 0.9$) between populations/species is important in reducing the frequency of negative cline widths.

#### S2: Descriptive cline fitting

Prior to cline fitting of SNP KASP data for the six colour loci, the allele frequencies in demes were collapsed to one-dimension, by using a linear transect through the approximate cline centre ($p = 0.5$ isocline of ROS1). To search for the optimal transect gradient we compared cline width and maximum log Likelihood (*logL*) values with a range of gradients and intercepts centred on the p = 0.5 isocline through the valley (see Fig S3 for example) at each of the loci.

Comparing the optimal transect across loci, there was no common transect direction (gradient) with a best fit across all loci (Fig S3). We found that a gradient of -0.345 and intercept at 6.9km generated the best compromise with highest likelihood (*logL; maxLL*) or within 2 *logL* of the maximum (maxLL) at two of the three loci (ROS1 and FLAVIA up; Fig S3). The third locus (FLAVIA down) was a considerably poorer cline fit at this transect, however comparing its best fitting transect cline parameters to the common well-fitting transect (at ROS1 and FLAVIA up) showed similar centres and widths regardless of the transect chosen. Therefore, from hereon we report cline parameters for the gradient of -0.345 and intercept at 6.9km (Fig S4).

#### S3: Clinal loci gene identification and enrichment

*Clines and gene classifications*

All major cline clusters on each chromosomes (top ranked cline proportions; Fig 4c) were identified from the annotation file corresponding to the reference genome version 3.5. The extracted gene sequences were formatted in FASTA and annotated using two primary bioinformatics tools. First, functional annotation was performed using the eggNOG-mapper v2, an efficient tool for fast functional annotation of novel sequences against the eggNOG database, leveraging orthology assignments, functional descriptions, and gene ontology (GO) terms (Huerta-Cepas et al., 2019). The eggNOG-mapper provides comprehensive insights into gene functions, significantly facilitating the understanding of gene roles within the genome (Huerta-Cepas et al., 2019). Simultaneously, the genes were subjected to similarity searches using the BLAST online tool (Basic Local Alignment Search Tool) against the NCBI non-redundant (nr) database. The BLAST results provided additional layers of functional evidence by identifying homologous sequences and potential gene functions based on sequence similarity. The functional information and gene hits obtained from both eggNOG-mapper and BLAST were manually curated to ensure accuracy and relevance. This manual curation process involved verifying the automated annotations, assessing the significance of BLAST hits, and reconciling discrepancies between different sources of functional evidence.

The location of all known genes known to influence flower colour was combined with genes identified around clinal loci. Genes were then categorized into functional groups, including the broad term ‘colour related gene’ to include all of those involved in the flavonol biosynthetic pathway or known to regulate the expression (intensity or distribution) of colour pigments across parts of the flower in *Antirrhinum majus*. Some of these genes have been confirmed to influence phenotypic variation between *pseudomajus* and *striatum* (e.g. MYB-related transcription factors *Rosea* and *Eluta*). However, others have been identified through genetic screens between wild type and mutant lines in *Antirrhinum majus,* but their importance for differences between these subspecies is unknown.

#### S4: Colour associations in hybrid zone plants and F2 crosses

We phenotyped and genotyped plants from the hybrid zone and controlled crosses, supported by RNAseq of extreme phenotypes to examine the effect of clinal loci on flower colour. In the hybrid zone, we randomly selected a subset of the larger KASP genotyped individuals (see above), photographs were taken on a black background with a colour standard. To perform more detailed colour quantification on a subset of plants, we stratified the random samples from the six major colour phenotypes (Magenta, Yellow, Pink, Weak Orange, Full Orange and White; following Whibley et al., (2006)), to ensure similar numbers of the main phenotype classes were available for genotype-phenotype associations.

Flower color measurements were taken in ImageJ (http://imagej.nih.gov/ij/). Images were white balanced using the macro `Chart_White_Balance` (https://imagejdocu.list.lu/plugin/color/chart_white_balance/start) which operated consistently across flowers with the color chart contained in the flower photos. To properly white balance the photos, a line was drawn from the whitest part of the white swath from the upper color chart to the darkest part of the dark swath of the upper color chart. The macro then operated and white-balanced the photo. Measurements were taken from each of six flower measurement location sites. These measurements were taken using a standard circle with an area of 5480 pixels. The circle was placed in six standard locations (Fig S15) using the plugin "RGB Measure" to obtain individual mean and standard deviation for the Red, Green, and Blue channels.

Hue was calculated using these three colour channels. We first convert all RGB measures to 0-1 scale, then calculate Hue depending on the maximum channel as (i) Red Max: Hue =[(G-B)/(max-min)]x60, (ii) Green Max: Hue= [2+(B-R)/(max-min)]x60, (iii) Blue Max: Hue= [4+[R-G]/(max-min)]x60. As a measure of colourfullness of each region in proportion to its brightness, we calculated Saturation (Sat) as Sat = [(max-min)/255]/[1-(2L-1)], where L= [0.5(max-min)/255] and max and min refer to the maximum and minimum value amongst Red, Green and Blue channels (rescaled between 0 and 1). We also calculated the standard deviation of the Grayscale (sd_GS), using the average of standard deviation values of Red, Green and Blue channels simply as GS_sd = (sd_R+sd_G+sd_B)/3. We next calculated intensity density (Int) averaged over the three channels as Int = (R+G+B)/3. Lastly, we converted the RGB colour scheme to HSV space. This was calculated first for Hue by rescaling to the 0-1 range, keeping Saturation (S) as for RGB and calculating Value (V) as the max intensity of the three channels.

To examine the association between genotypes at *RUBIA* and colour scores we use linear regression. First we linearise Hue scores (to account for $360^0$ wrparound effects of Hue scale), by by calculating the median Hue, and using the linear distance to the median to convert to linear scale. We then fit a linear model which included the effects of each RUB and ROS1 and their interaction effects (RUB + ROS + ROS:RUB) using R (e.g. lm(H_4 ~ RUB+ ROS1 + RUB:ROS1, data = colourData). These models were repeated for each colour region, for Hue, Saturation and Value scores. An ANOVA was used to compare different models, showing that linear regressions with interaction effects fitted better than with no interactions effects ($p < 0.05$).

#### S5: RNA seq and differential gene expression analysis

RNA was extracted from corolla tissue from snapdragon flowers for three biological replicates of a representative *A. majus ssp pseudomajus* (magenta flowers) and *A. majus ssp striatum* (yellow)(See Bradley et al,. 2017). We used the annotated *Antirrhinum* reference genome v3.5 and added "decoy-aware" index for the mapping. For each of the samples we next used Salmon to align and quantify transcript abundance, including flags to increase the stringency of the mappings (--validateMappings) and to learn and apply corrections for GC bias and primer bias (--gcBias --seqBias). We then performed differential gene expression analyses using DESeq2 (Bioconductor) with the lfcShrink function to compensate for inflated log2fold changes in genes that have low counts (after filtering out genes with < 10 reads). As a reference point, we checked known colour genes in the top clusters on Chr 6 (ROS/EL) and Chr 2 (FLA) and found as expected high differential gene expression.

### SI Figures

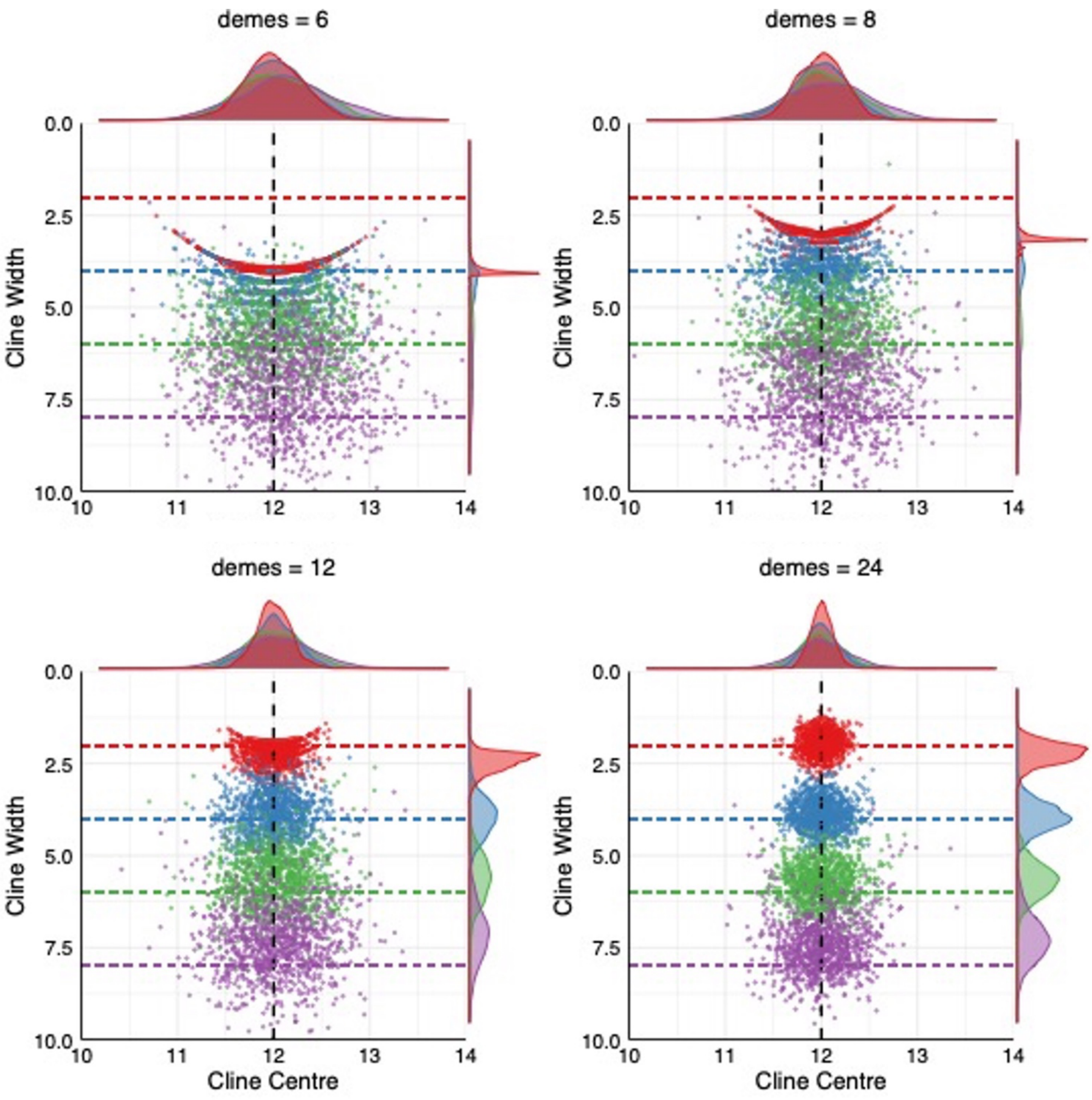

**Fig S1. Simulations of FastClines estimates with varying cline widths and number of demes for loci with fixed differences and high sequencing depth.**

Panels show estimates of cline width and centres from 10,000 simulations of sampling of individuals (alleles from demes) from symmetric geographic sigmoid clines for (a) 6 demes, (b) 8 demes, (c) 10 demes, (d) 12 demes. Within each panel, simulated clines were run at each of four cline width values (2, 4, 6 and 8 km) and cline centre fixed at 12km. For all simulation runs sequencing depth was set to 100, $F_{ST}$= 0.01, $F$ = 0.

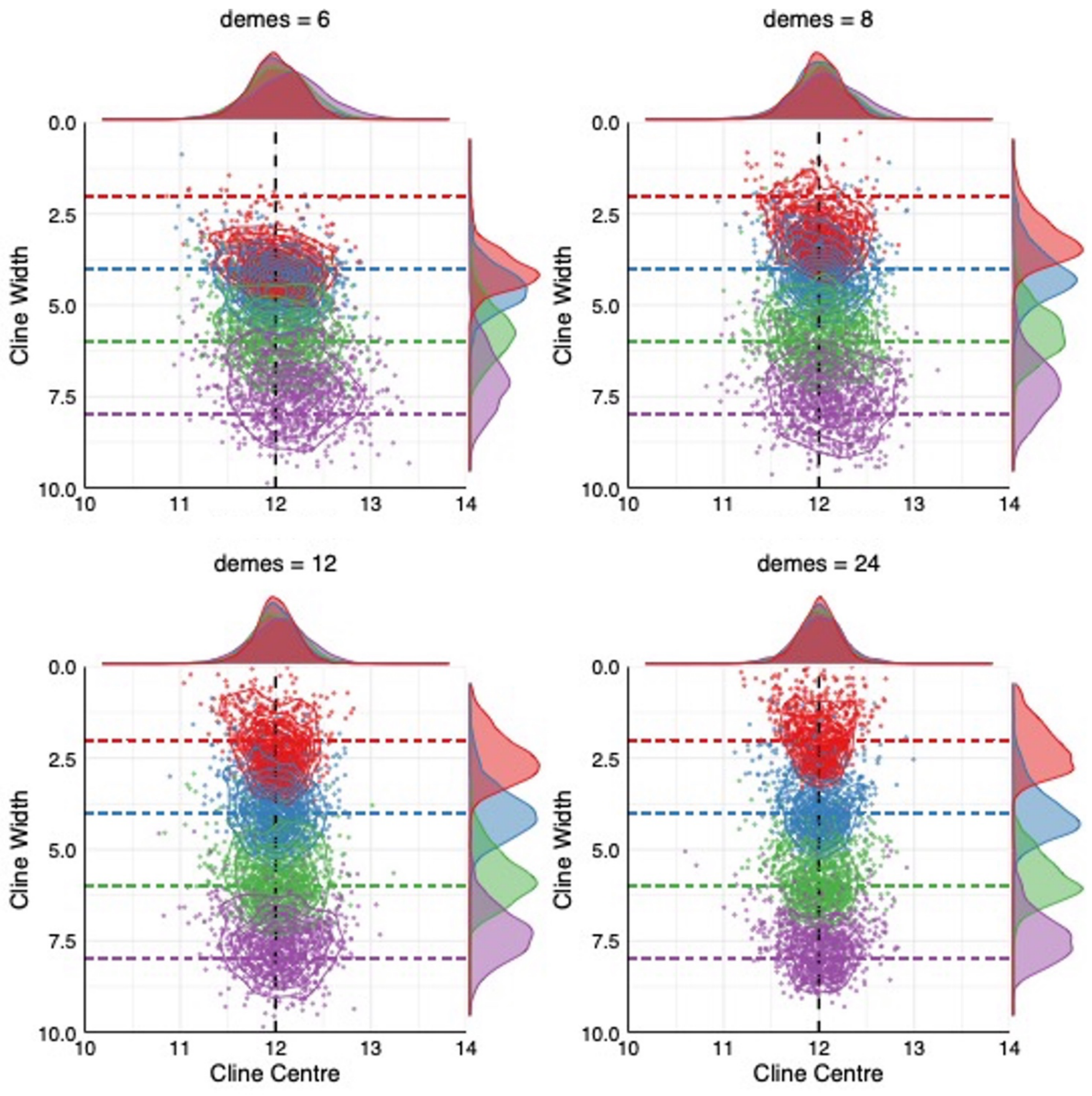

**Fig S2. Simulations of FastClines estimates of cline width and centre for loci with AFD = 0.95.**

Panels show estimates of cline width and centres from 10,000 simulations of sampling of individuals (alleles from demes) from symmetric geographic sigmoid clines for (a) 6 demes, (b) 8 demes, (c) 10 demes, (d) 12 demes. Within each panel, simulated clines were run at each of four cline width values (2, 4, 6 and 8 km) and cline centre fixed at 12km. For all simulation runs sequencing depth was set to 100, $F_{ST}$= 0.01, $F$ = 0.

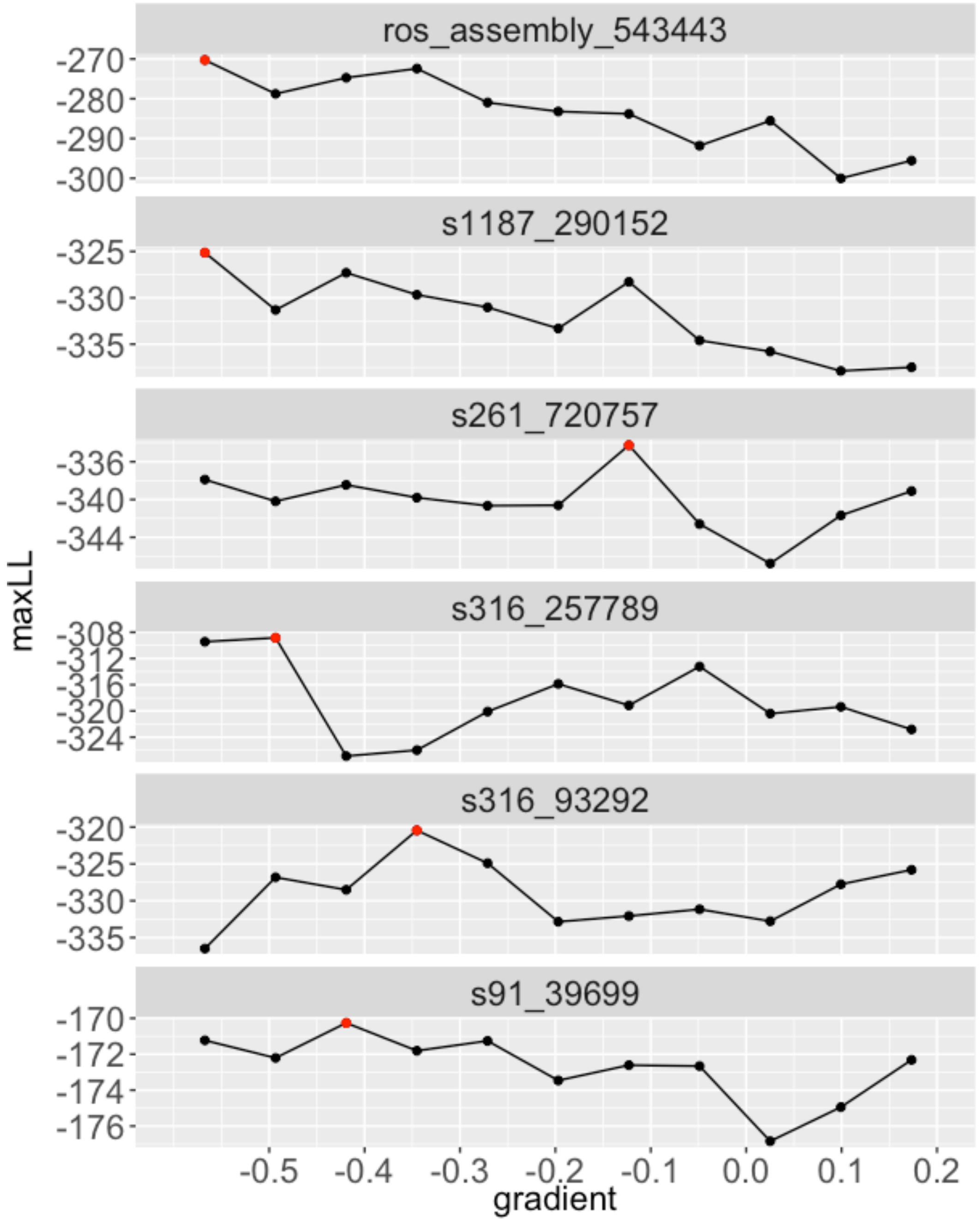

**Fig S3. Maximum log Liklihood (maxLL) for each gradient parameter search at each transect directions**

Transect (as gradients) through the hybrid zone at SNP markers at each of six key flower colour loci (ros_assembly_543443 = *ROS*, s1187_290152 = *CRE*, s261_720757 = *RUB*, s316_257789 = *FLA1*, s316_93292 = *FLA2*, s91_39699 = *SULF*). The highest maxLL indicated with red filled circle.

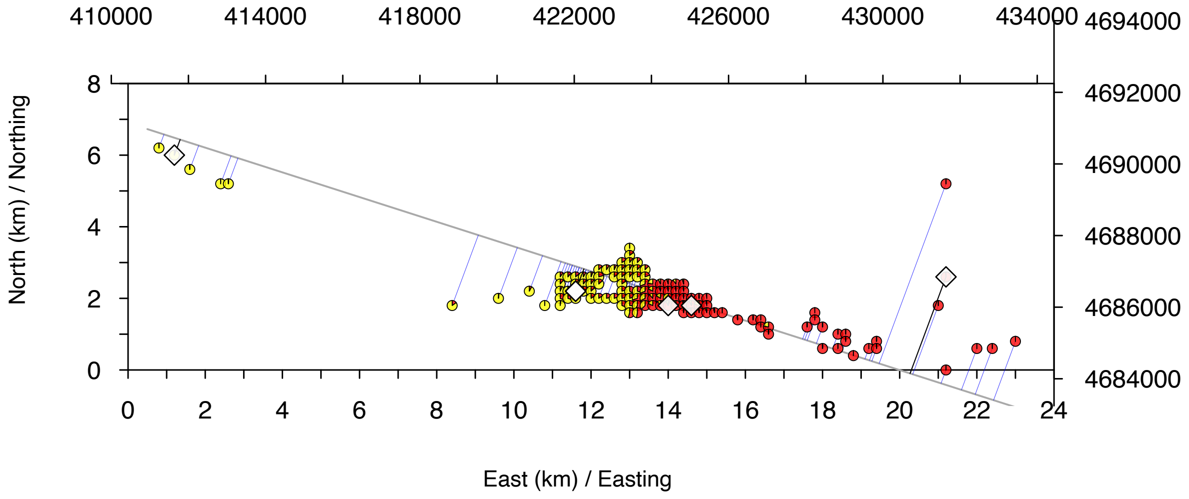

**Fig S4. Allele frequencies from KASP SNP genotyping at the *ROS1* locus in 200m demes along the hybrid zone transect.**

Each pie chart indicates allele frequency proportions as yellow = *A. majus majus var. striatum*, red = *A. majus majus var. pseudomajus*. Each deme is collapsed to a geographic distance along the transect (black line) that is perpendicular (blue line) with the deme. Location of six whole genome PoolSeq demes shown as white diamonds. Distance along the hybrid zone show in kilometres and in Easting and Northings. Linear transect show is the optimal line chosen for cline fits with a gradient -0.345 and intercept at 6.9km.

#### Chromosome 2: *FLA*

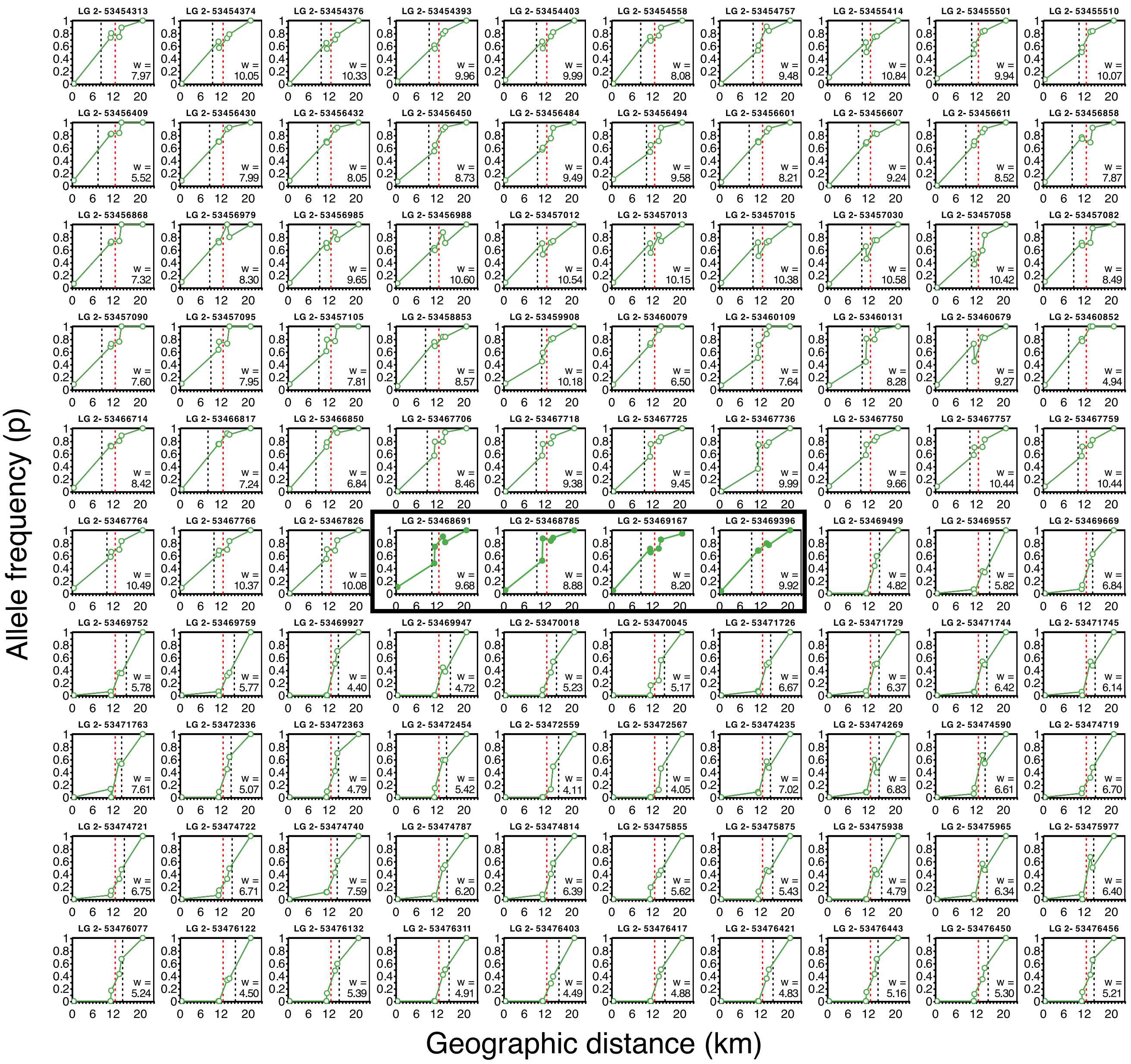

**Fig S5. Allele frequencies for clinal loci around the *FLA* gene on Chromosome 2.**

An example of clinal loci located within *FLA* coding sequences (plots within black box) and upstream (plots left of black box) and downstream (plots right of black box). For each plot, the header indicates the Chromsome number followed by the genomic position of the locus. Phenotypic cline centre (red dashed line), FastCline estimate of cline centre (black dashed line) and width (w) indicated within each plot.

#### Chromosome 2: >300kb *FLA*

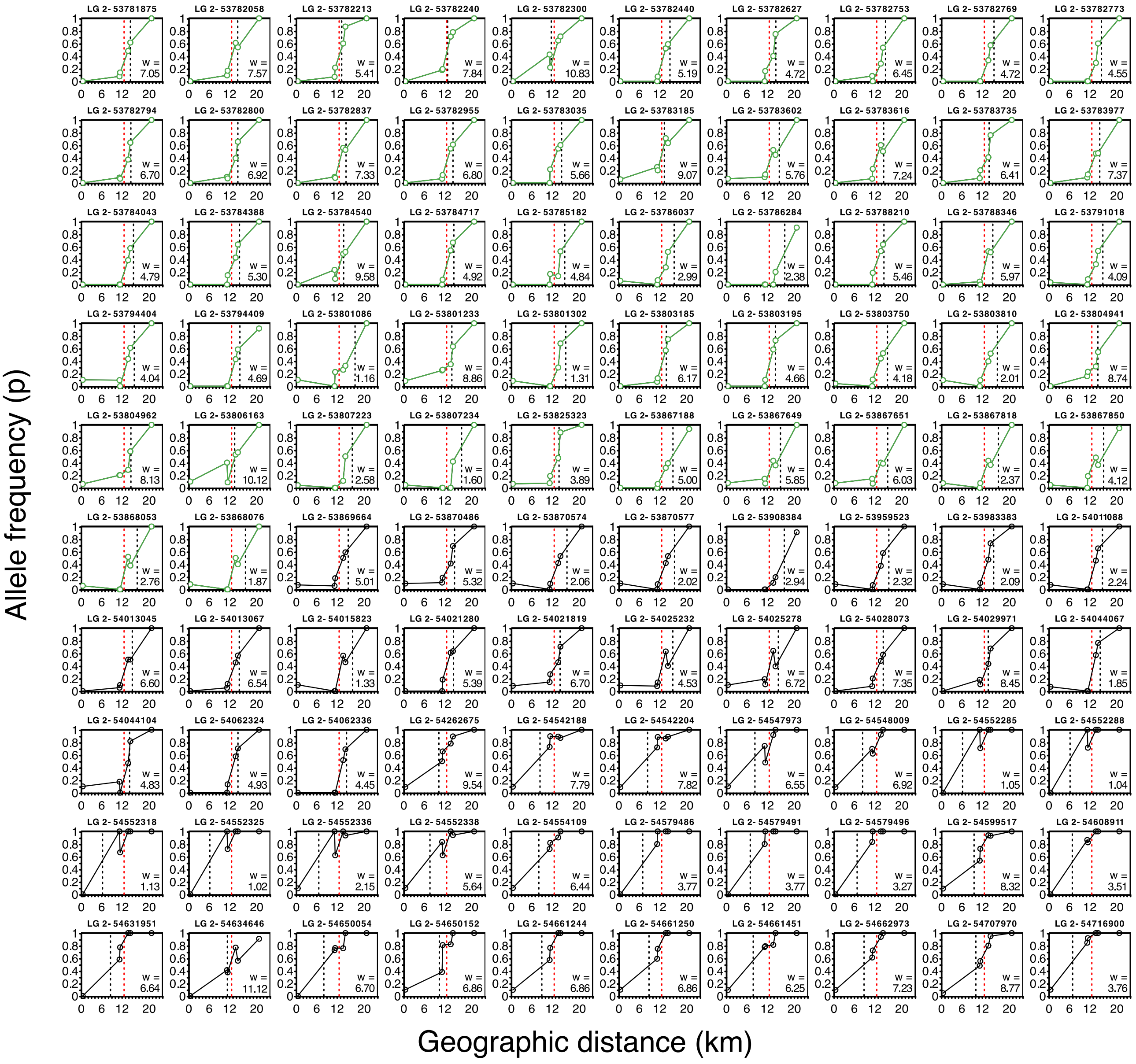

**Fig S6. Allele frequencies for clinal loci downstream of *FLA* gene (>300kb) on Chromosome 2.**

An example of clinal loci located further downstream of *FLA* gene (>300kb) illustrating the cline centres shifted towards the yellow flank of hybrid zone. Clinal loci located <300kb (allele frequencies in green lines and green open circles) and clinal loci located >300kb from *FLA* gene (black lines and open black circles). For each plot, the header indicates the Chromsome number followed by the genomic position of the locus. Phenotypic cline centre (red dashed line), FastCline estimate of cline centre (black dashed line) and width (w) indicated within each plot.

#### Chromosome 1: *CRE*

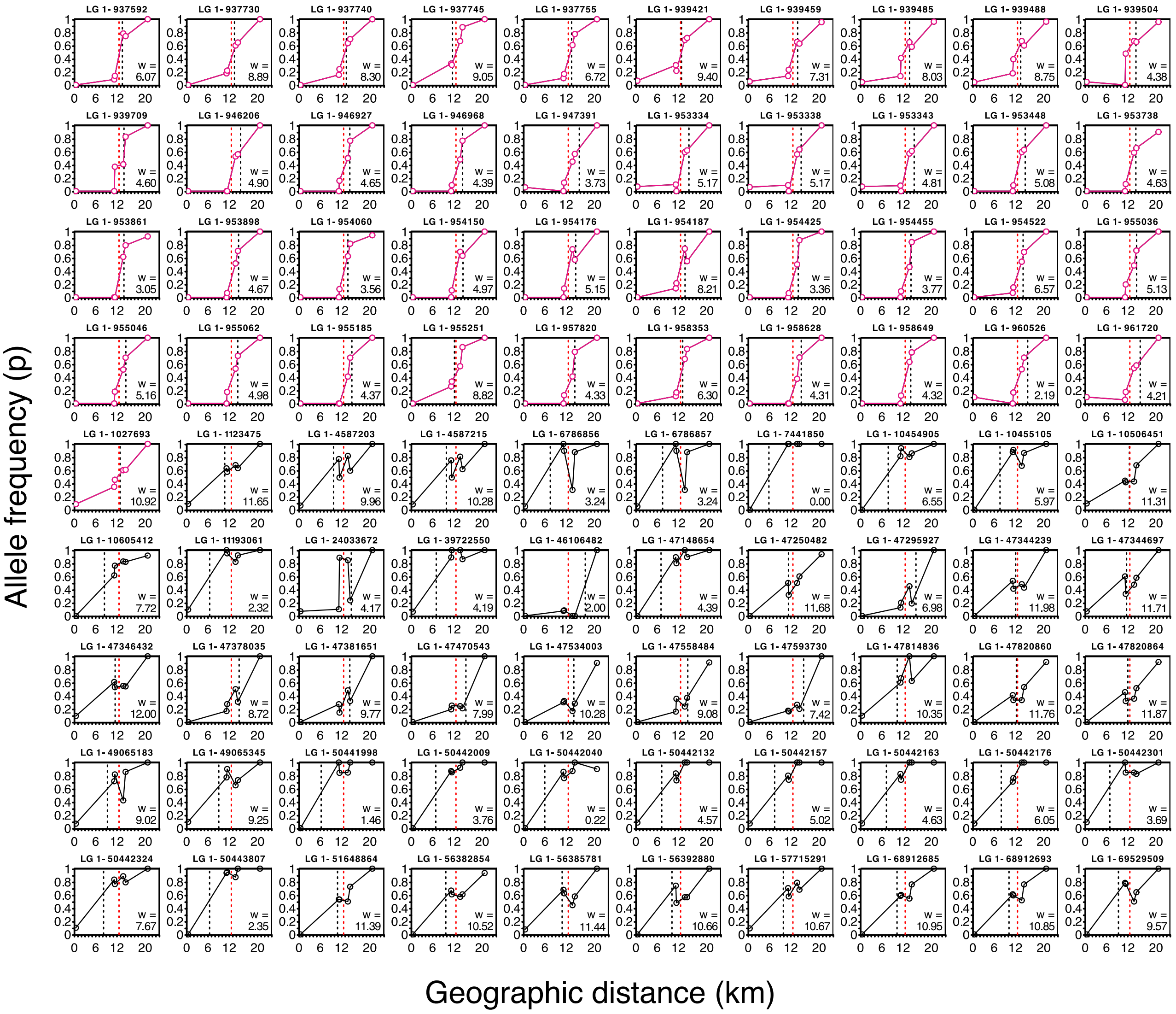

**Fig S7. Allele frequencies for clinal loci around the *CRE* gene on Chromosome 1.**

All clinal loci on Chromosome 1, including loci located within 100kb of *CRE* coding sequences (plots with allele frequencies with pink lines and circles) and 'background' loci > 100kb distant (plots with allele frequencies with black lines and circles). For each plot, the header indicates the Chromsome number followed by the genomic position of the locus. Phenotypic cline centre (red dashed line), FastCline estimate of cline centre (black dashed line) and width (w) indicated within each plot.

#### Chromosome 4: *SULF*

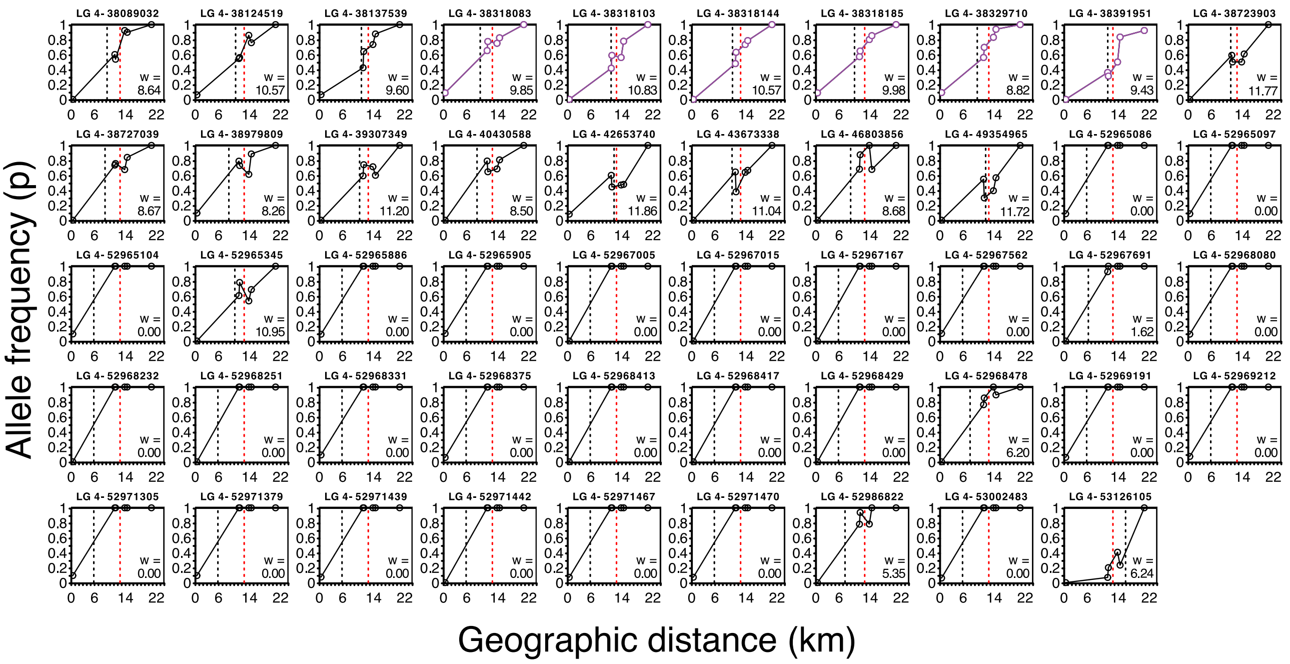

**Fig S8. Allele frequencies for clinal loci around the *SULF* gene on Chromosome 4.**

Clinal loci located <100kb from SULF region (plots with purple lines and circles for allele frequencies) and >100kb (plots with black lines and circles for allele frequencies). For each plot, the header indicates the Chromsome number followed by the genomic position of the locus. Phenotypic cline centre (red dashed line), FastCline estimate of cline centre (black dashed line) and width (w) indicated within each plot.

#### Chromosome 5: *RUB*

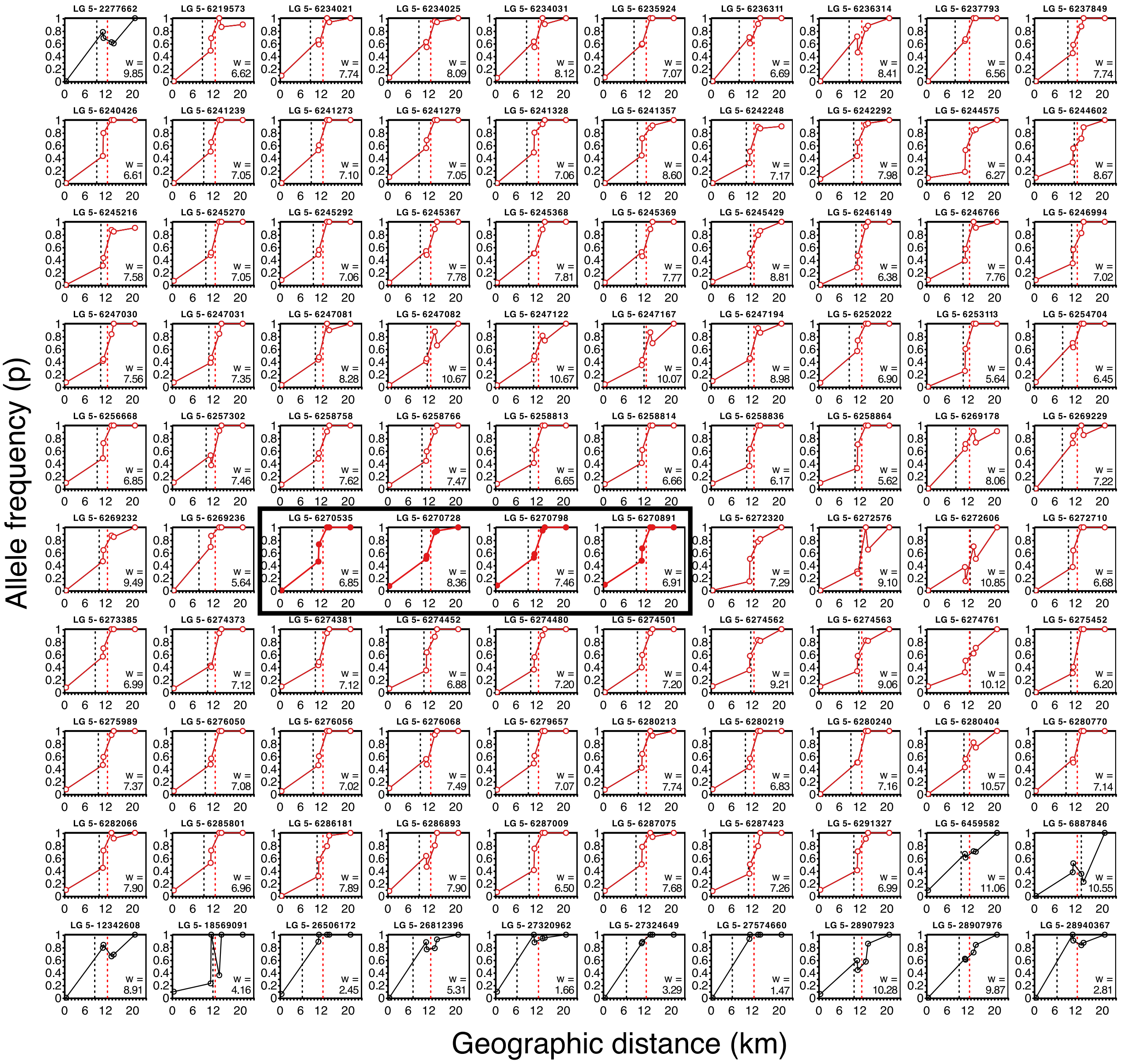

**Fig S9. Allele frequencies for clinal loci around the *RUB* gene on Chromosome 5.**

Clinal loci located within RUB coding sequences (plots within black box) and upstream (plots left of black box) and downstream (plots right of black box). For each plot, the header indicates the Chromsome number followed by the genomic position of the locus. Phenotypic cline centre (red dashed line), FastCline estimate of cline centre (black dashed line) and width (w) indicated within each plot.

#### Chromosome 6: *ROS/EL*

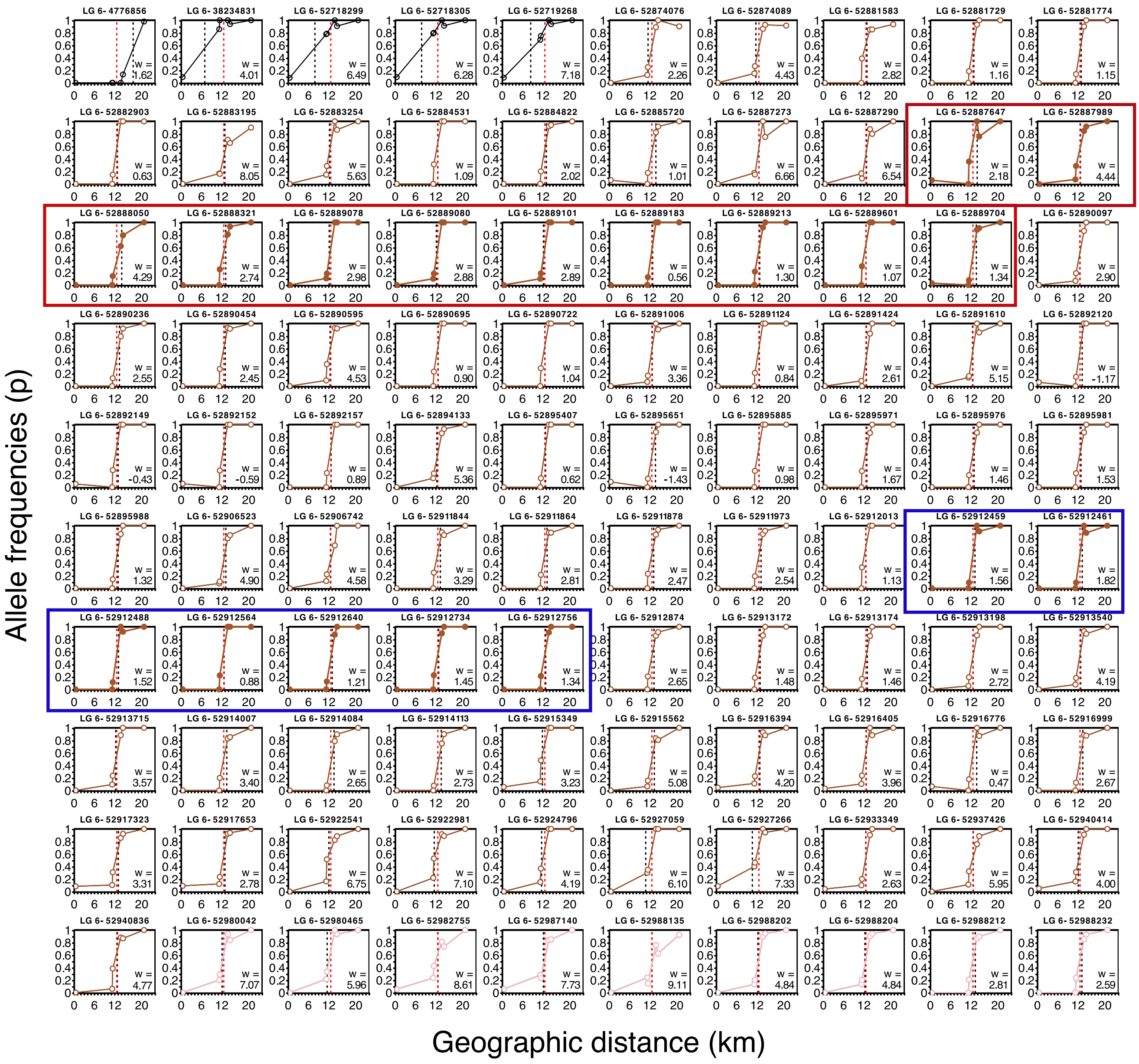

**Fig S10. Allele frequencies for clinal loci around the *ROS*/*EL* genes on Chromosome 6.**

An example of clinal loci located within *ROS* coding sequences (plots within red boxes), within *EL* gene (blue boxes) and upstream and downstream of both genes (plots left and right of coloured boxes). For each plot, the header indicates the Chromsome number followed by the genomic position of the locus. Phenotypic cline centre (red dashed line), FastCline estimate of cline centre (black dashed line) and width (w) indicated within each plot.

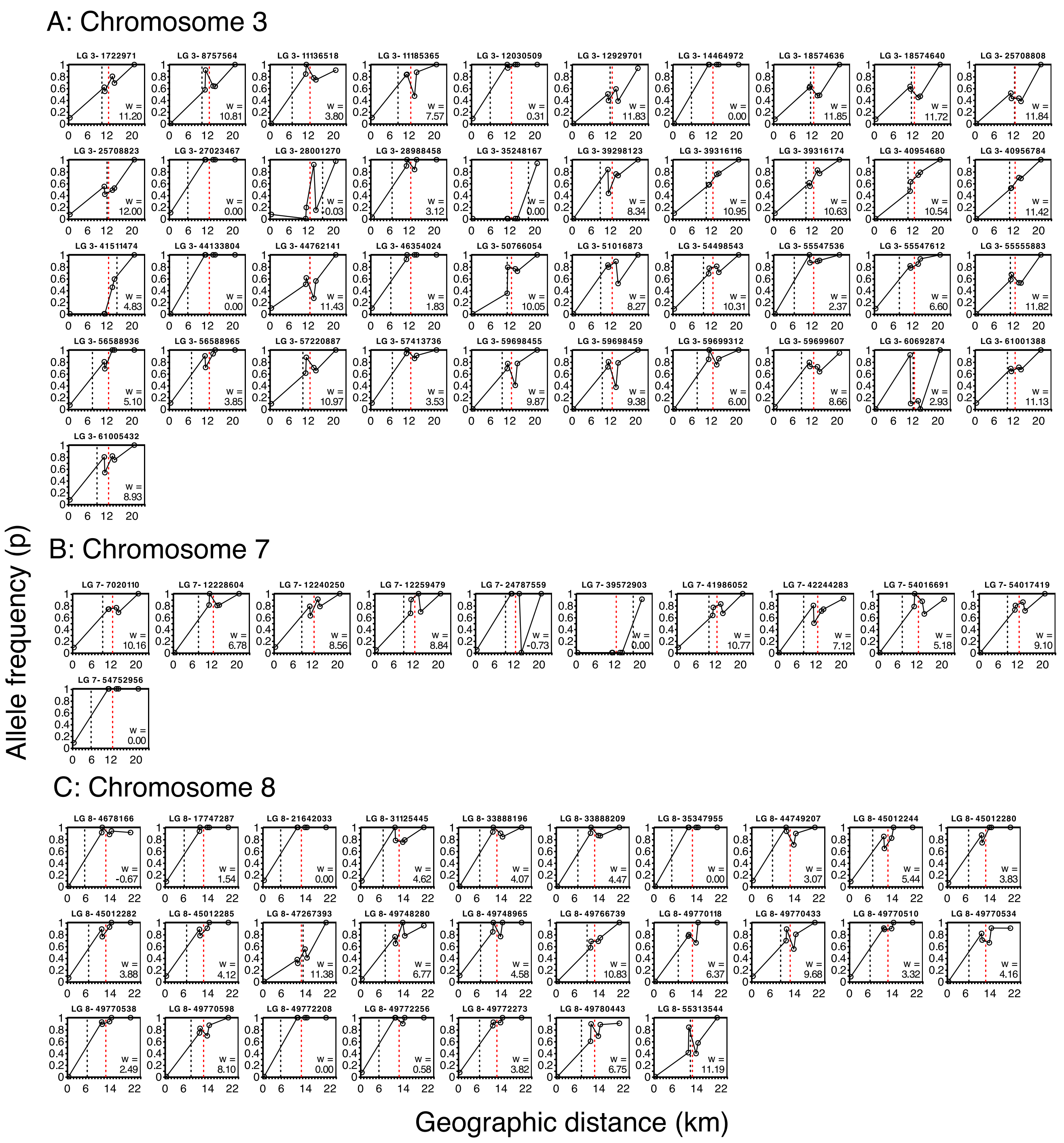

**Fig S11. Allele frequencies for clinal loci around on Chromosome 3, 7, and 8.**

All clinal loci located on (A) Chromsome 3, (B) Chromosome 7, and (C) Chromosome 8. For each plot, the header indicates the Chromosome number followed by the genomic position of the locus. Phenotypic cline centre (red dashed line), FastCline estimate of cline centre (black dashed line) and width (w) indicated within each plot.

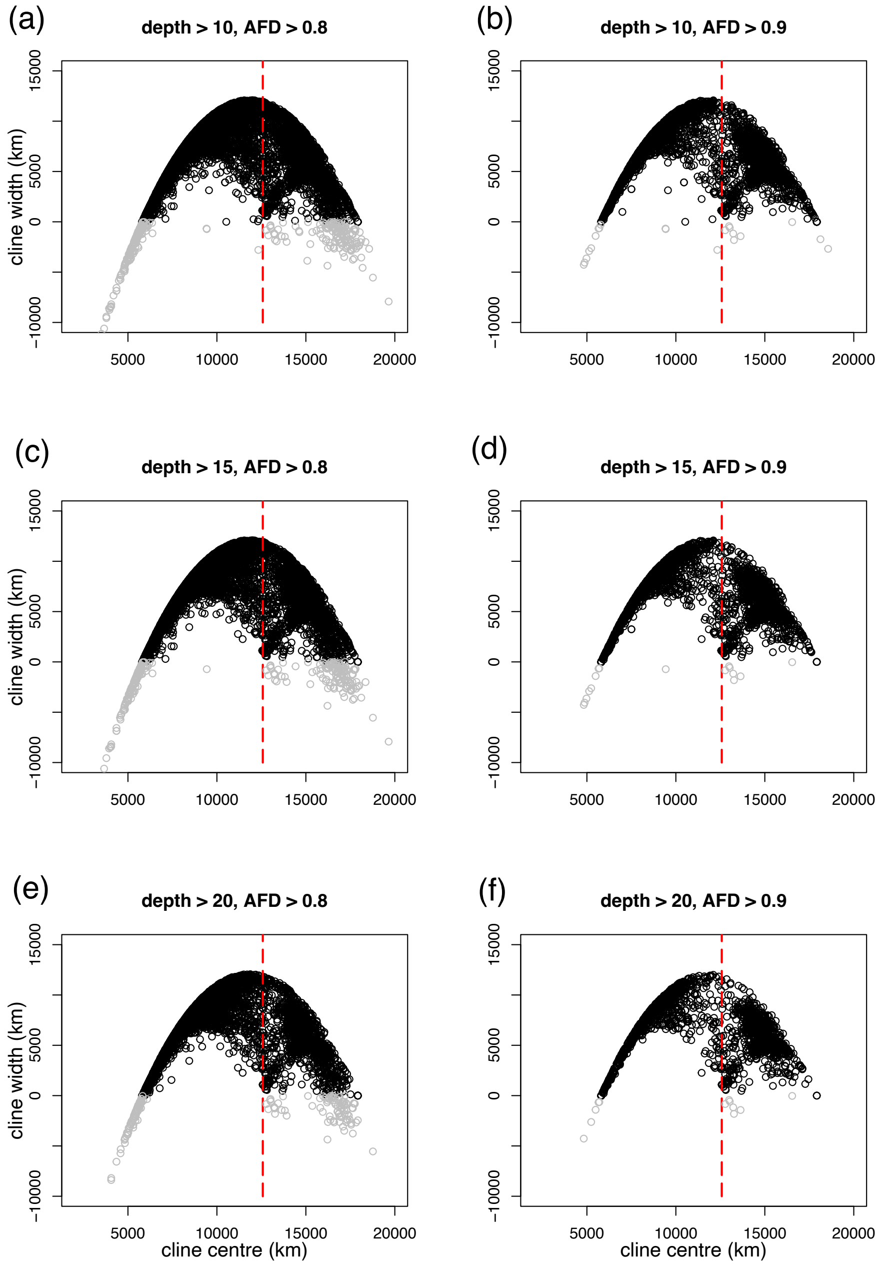

#### Fig S12. Cline width and centre for different filtering for loci

With a minimum of (a) depth 10 (in 5 of the 6 pools) and Alelle Frequency Difference (AFD) of $\Delta p_{1,6} \geq 0.80$ (between pool 1 and 6), (b) depth 10 and AFD $\Delta p_{1,6} \geq 0.90$, (c) depth 15 and AFD $\Delta p_{1,6} \geq 0.80$, (d) depth 15 and AFD $\Delta p_{1,6} \geq 0.90$, (e) depth 20 and AFD $\Delta p_{1,6} \geq 0.80$, (f) depth 20 and AFD $\Delta p_{1,6} \geq 0.90$. The SNPs include loci with positive cline widths (black) and negative cline widths (grey). Dashed red line indicate phenotypic cline centre.

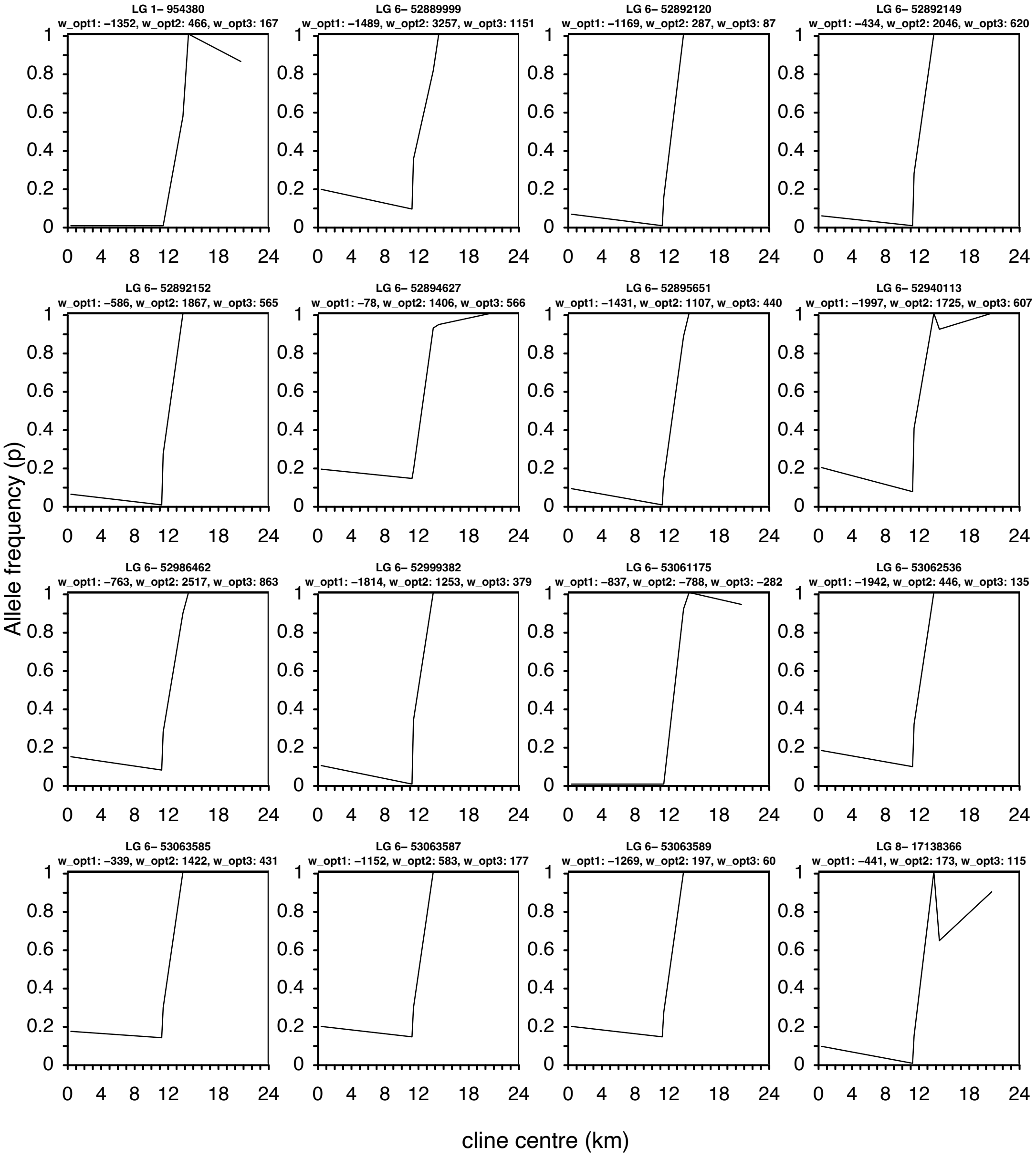

**Fig S13. Allele frequencies across the transect for 16 loci with negative cline widths** with depth filtering of 15, Deme Span 1 and Alelle Frequency Difference (AFD) of $\Delta p_{1,6} \geq 0.80$.

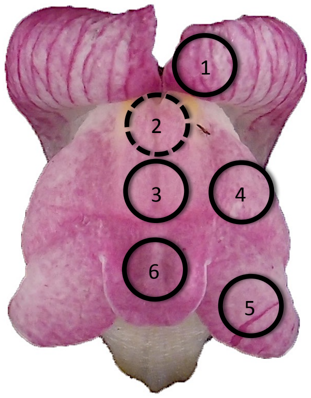

**Fig S14. An example of the six regions of the front view of an *Antirrhinum* flower used to quantify patterns of floral pigmentation.**

The circle 2 is indicated as a dashed line because these measurements were not used in the analyses. This was due to some flowers having ventral petals that drooped over the 2nd measurement, making the spacing inaccurate and not comparable between flowers.

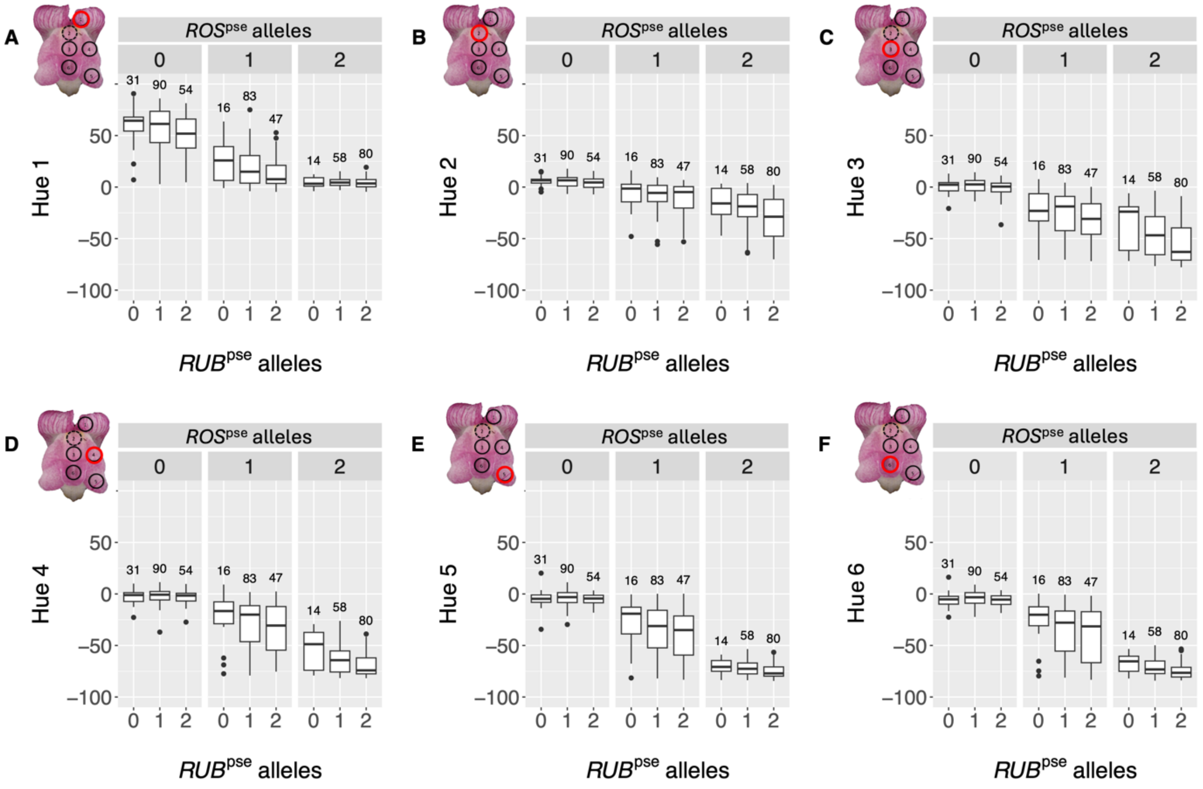

**Fig S15. Summary of HSV Hue scores of *Antirrhinum* flowers for *ROS* and *RUB* haplotypes from the hybrid zone.**

Hue (in HSV colouor space) in six regions of the flower (a – f) for 473 plants from the hybrid zone. For each panel, plants are grouped haplotypes, first via facets which group plants by the number of copies of *ROS* alleles from *A. m. m. pseudomajus* (*ROS*$^{pse}$) and secondly by the number of *RUB* alleles from *A. m. m. pseudomajus* (*RUB*$^{pse}$) along x-axis. The numbers above each box indicate sample size . Insets of flower images indicate the focus region of the Hue measurements with a red circle (see Fig S15 for details).

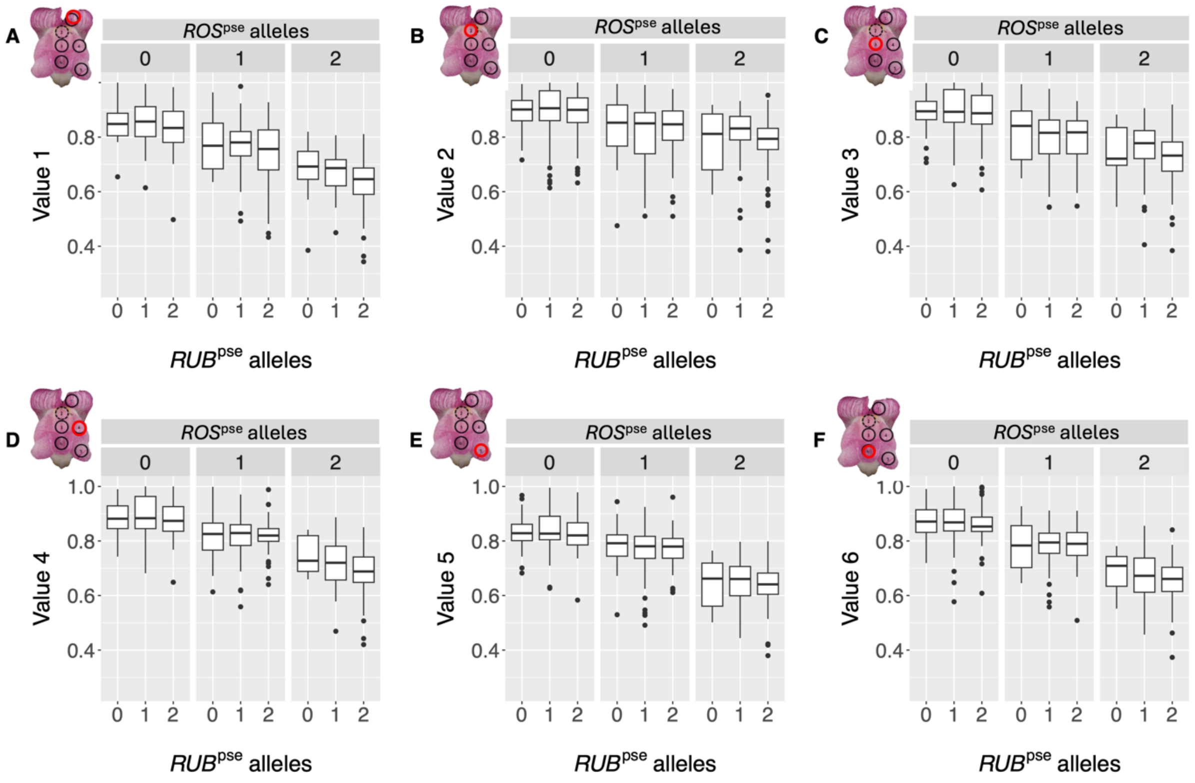

**Fig S16. Summary of HSV Saturation scores of *Antirrhinum* flowers for *ROS* and *RUB* haplotypes from the hybrid zone.**

Saturation (in HSV colour space) in six regions of the flower (a – f) for 473 plants from the hybrid zone. For each panel, plants are grouped haplotypes, first via facets which group plants by the number of copies of *ROS* alleles from *A. m. m. pseudomajus* (*ROS$^{pse}$*) and secondly by the number of *RUB* alleles from *A. m. m. pseudomajus* (*RUB$^{pse}$*) along x-axis. The numbers above each box indicate sample size. Insets of flower images indicate the focus region of the Saturation measurements with a red circle.

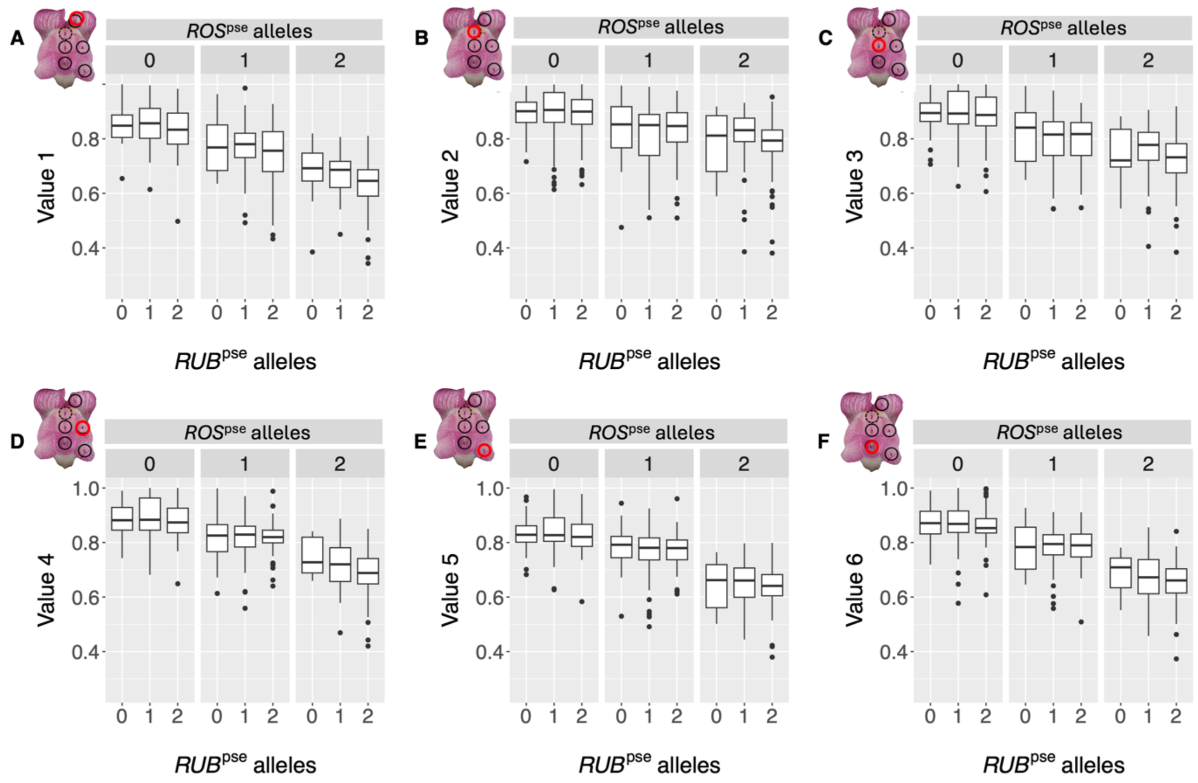

**Fig S17. Summary of HSV Value scores of *Antirrhinum* flowers for *ROS* and *RUB* haplotypes from the hybrid zone.**

Saturation (in HSV colour space) in six regions of the flower (a – f) for 473 plants from the hybrid zone. For each panel, plants are grouped haplotypes, first via facets which group plants by the number of copies of *ROS* alleles from *A. m. m. pseudomajus* (*ROS$^{pse}$*) and secondly by the number of *RUB* alleles from *A. m. m. pseudomajus* (*RUB$^{pse}$*) along x-axis. The numbers above each box indicate sample size. Insets of flower images indicate the focus region of the Saturation measurements with a red circle.

### SI Tables

#### Table S1. Geographic locations of whole genome pools across *Antirrhinum* hybrid zone

Details of deme locations at *Antirrhinum majus subspecies majus* hybrid zone at Planoles. For the first deme, position zero set from the first deme from the larger KASP SNP genotyping in relation to whole genome pools (poolSeq) on the yellow side with *A. m. m var striatum* (YP4, YP1, YP2) and magenta side *A. m. m var pseudomajus* (MP2, MP4, MP11). Distance along the transect (Fig 1), deme span ($d_i$), number of samples included in poolSeq, and mean depth and standard deviation (in parentheses).

| deme/pool | Easting | Northing | distance along transect (km) | deme span | number of samples | mean depth (SD) |
|---|---|---|---|---|---|---|
| KASP Start | 411310.104 | 4690519.39 | 0 | - | - | |
| YP4 | 411660.104 | 4690319.39 | 0.38 | 6000 | 50 | 53.3 (21.7) |
| YP1 | 421960.104 | 4686519.39 | 11.23 | 5000 | 50 | 24.2 (6.8) |
| YP2 | 422110.104 | 4686369.39 | 11.40 | 1500 | 50 | 22.6 (6.3) |
| MP2 | 424460.104 | 4686069.39 | 13.77 | 1400 | 50 | 22.5 (6.2) |
| MP4 | 425110.104 | 4685969.39 | 14.42 | 3500 | 50 | 25.8 (7.2) |
| MP11 | 431660.104 | 4686869.39 | 20.67 | 6000 | 50 | 39.3 (15.4) |

#### Table S2. KASP SNP genotype marker details

Details of the KASP marker design including locus marker name (with LGC), colour loci, the colour pigment locus is associated with, the chromosome (Chr), position on chromosome and the sequence for the oligo design in IUCN code. The biallelic SNP is indicated (e.g. [A/C]).

| Locus marker name | colour loci | colour pigment | Chr | position | Oligo |
|---|---|---|---|---|---|
| s1187_290152 | *CRE* | yellow | 1 | 955185 | GTCTCACATTGTTGACAAAATCCAAGTCGTGACTTGGGAGGAAGAATAAT[A/C]AAAATGGTCATGRTMRAGTCACTTTYCCCTTAATCACCAAAATAGAAAAA |
| s316_93292 | *FLA 1* | yellow | 2 | 53594773 | AGATTTGCTATMTGGTATTGGAGTCGCTGGAAGATAAAGAAAGTACGCCA[A/G]TCGAGCTAAACCTCCAGCTTCCTGGTTATGATGGAAACAAAGAACATGAA |
| s316_257789 | *FLA 2* | yellow | 2 | | TGTATGTGGCAGCTTCACATTATACACAATTGCATGCAGACGAACCAATA[A/G]CCAGGGGCGTATCYARGAATTTTATCTGGGGAGGGCTAAYCTAATGAATA |
| s91_39699 | *SULF* | yellow | 4 | 38316531 | GGTAATCAAGCACATAAAAYATTAATAACAAGATTAYGAAATYARATCAA[C/T]TGGTCCACACAYATAAATCATAACACAACAATCATAACATAACCAGATTC |
| s261_720757 | *RUB* | magenta | 5 | 6286923 | CAAAGTAYGMCATTTGCACCYATTCATTTGAGAGCTCAACGATCGAATAT[C/T]GATCATAACCTCGATTTGGATCGTGCTCTTTCCCAGTTCCRTCTTGTCTC |
| ros_assembly_543443 | *ROS 1* | magenta | 6 | | GAAACTMAAAAATTMAAGATAAATTTGCTCGTGTCAATARTAGTGAAAACTAT[A/G]CWTCAATTAGTTATTGAAAATGWAACCTGTCTATTTCTATAAGTGTTTAGCG |
| ros_assembly_715015 | *EL* | magenta | 6 | 53061094 | AGAYGTGAATTCCAATGGYTCTTCACTTCATCATCCRTTCGSCCCGGYAA[A/G]CGTCCCGCAATTAGAGACCACCTAACAGTMAAGAAAGTGAATTAATAYAA |

#### Table S3. MLE cline parameters for best fitting model of KASP SNP genotypes

The best fitting cline parameters for polymorphic sigmoid cline fits with simulated annealing. Estimates at each of the colour linked loci (Locus marker name) associated with each Colour loci, the chromosome (Chr), followed by the four main parameters – cline centre, cline width, allele frequencies on the left (west) of transect for *A. m. m. var. striatum* (p0) and the right (east) of transect for *A. m. m. var. pseudomajus* and their 95% confidence intervals (in parentheses).

| **Locus marker name** | **Colour loci** | **Chr** | **centre** | **width** | **p0** | **p1** |
|---|---|---|---|---|---|---|
| s1187_290152 | *CRE* | 1 | 13.9 (13.7 - 14) | 3.1 (2.7 – 3.4) | 0.05 (0.04 - 0.06) | 0.91 (0.90 - 0.92) |
| s316_93292 | *FLA* 1 | 2 | 13.9 (13.8 – 14.2) | 3.0 (2.6 – 3.6) | 0.03 (0.01 - 0.04) | 0.94 (0.91 - 0.97) |
| s316_257789 | *FLA* 2 | 2 | 10.5 (10.2 – 10.9) | 9.1 (8.5 – 10.1) | 0.11 (0.0 - 0.18) | 0.99 (0.99 - 0.99) |
| s91_39699 | *SULF* | 4 | 13.5 (13.5 – 13.6) | 934 (0.9 – 12.5) | 0.32 (0.32 - 0.32) | 0.93 (0.93 - 0.94) |
| s261_720757 | *RUB* | 5 | 11.9 (11.5 – 12.4) | 7.6 (6.0 – 9.1) | 0.13 (0.06 - 0.23) | 0.95 (0.90 - 0.97) |
| ros_assembly 543443 | *ROS1* | 6 | 13.2 (13.1 – 13.3) | 876 (0.8 – 10.2) | 0.11 (0.09 - 0.12) | 0.99 (0.98 - 0.99) |

**Table S4. Test for greater clustering of divergent loci showing clines on each LG (Chromosome) than expected by chance.**

LG: linkage group (chromosome). Obs. mean pairwise distance: the mean pairwise distance between clinal windows in Mbp. The mean pairwise distance in Mbp calculated from the 99,999 random permutations. The p-values show the probability of obtaining the observed mean pairwise distance by chance.

| LG | *n* clines | Obs. mean pairwise distance (Mbp) | Mean perm pairwise distance (Mbp) | p-value |
|---|---|---|---|---|
| 1 | 21 | 25.3 | 24.2 | 0.340 |
| 2 | 170 | 12.3 | 25.6 | 1e-5 |
| 3 | 22 | 22.1 | 21.0 | 0.295 |
| 4 | 49 | 14.9 | 18.8 | 0.001 |
| 5 | 28 | 20.8 | 24.0 | 0.076 |
| 6 | 22 | 5.7 | 18.6 | 1e-5 |
| 7 | 5 | 25.3 | 18.7 | 0.095 |
| 8 | 6 | 21.8 | 18.9 | 0.288 |

**Table S5. Test for enriched overlap of clinal windows and $F_{ST}$ outliers.**

Pops: the populations that $F_{ST}$ was calculated between. ΔP: the allele frequency cut-off used in the *fastclines* analysis. n clines: the number of clines detected in the *fastcline*s analysis. Observed overlaps: the number of clinal windows that were also $F_{ST}$ outliers. Mean perm overlaps: the mean number of overlaps, calculated from the 99,999 random permutations. The p-values show the probability of obtaining the observed number of overlaps by chance

| Pops | Δ*P* | *n* clines | Observed overlaps | Mean perm overlaps | p-value |
|---|---|---|---|---|---|
| YP1-MP6 | 0.8 | 1311 | 753 (57.4%) | 65.53 (5.0%) | 1e-5 |
| YP1-MP6 | 0.9 | 323 | 225 (69.7%) | 16.14 (5.0%) | 1e-5 |
| YP1-MP6 | 1.0 | 136 | 111 (81.6%) | 6.79 (5.0%) | 1e-5 |
| YP3-MP4 | 0.8 | 1311 | 167 (12.7%) | 65.32 (5.0%) | 1e-5 |
| YP3-MP4 | 0.9 | 323 | 89 (27.6) | 16.06 (5.0%) | 1e-5 |
| YP3-MP4 | 0.8 | 136 | 57 (41.9%) | 6.75 (5.0%) | 1e-5 |

**Table S6. Test for differences in $\pi$, $d_{xy}$ and $F_{ST}$ between clinal and non-clinal windows between population pairs.**

DP: depth cut-off for including a site in the dataset. Pop1 and Pop2: IDs of the populations being compared. ΔP: the allele frequency cut-off used in the *fastclines* analysis. The absolute value of the observed difference between the mean clinal and non-clinal values loci are given. The p-values show the probability of obtaining the observed difference by chance, determined using a permutation test (99,999 random permutations).

| | | | | Obs. difference between means | | | | p-value | | | |
|---|---|---|---|---|---|---|---|---|---|---|---|
| DP | Pop1 | Pop2 | $\Delta P$ | $\pi_{pop1}$ | $\pi_{pop2}$ | $d_{xy}$ | $F_{ST}$ | $\pi_{pop1}$ | $\pi_{pop2}$ | $d_{xy}$ | $F_{ST}$ |
| 15 | YP1 | MP6 | 0.8 | 0.0004 | 0.0014 | 0.0024 | 0.1345 | 1.7e-3 | 1e-5 | 1e-5 | 1e-5 |
| 15 | YP1 | MP6 | 0.9 | 0.0012 | 0.0018 | 0.0041 | 0.2006 | 2e-5 | 1e-5 | 1e-5 | 1e-5 |
| 15 | YP3 | MP4 | 0.8 | 0.0005 | 0.0005 | 0.0004 | 0.0076 | 3.9e-4 | 1.7e-3 | 0.018 | 1e-5 |
| 15 | YP3 | MP4 | 0.9 | 0.0005 | 0.0002 | 9.7e-5 | 0.0231 | 0.052 | 0.527 | 0.737 | 1e-5 |
| 20 | YP1 | MP6 | 0.8 | 0.0015 | 0.0032 | 0.0019 | 0.1836 | 1e-5 | 1e-5 | 1e-5 | 1e-5 |
| 20 | YP1 | MP6 | 0.9 | 0.0024 | 0.0031 | 0.0044 | 0.2654 | 1e-5 | 1e-5 | 1e-5 | 1e-5 |
| 20 | YP3 | MP4 | 0.8 | 0.0022 | 0.0020 | 0.0020 | 0.0109 | 1e-5 | 1e-5 | 1e-5 | 1e-5 |
| 20 | YP3 | MP4 | 0.9 | 0.0017 | 0.0011 | 0.0008 | 0.0334 | 2e-5 | 5.1e-4 | 0.0008 | 1e-5 |

#### Table S7. Linear regressions of genotype variation at *ROS* and *RUB* loci and quantitative flower colour scores for Hue in HSV colour space.

Overall model (intercept), individual loci (RUB, ROS1) and their interaction (RUB:ROS1) effects, standard error, p-value ($p < 0.05$ in bold) and Model (the six floral regions in Fig S15)

| term | estimate | std.error | statistic | p.value | Model |
|---|---|---|---|---|---|
| (Intercept) | 47.95976 | 14.39679 | 3.331281 | **0.000933** | Hue 1 |
| RUB | 29.91842 | 10.79177 | 2.772336 | **0.005787** | Hue 1 |
| ROS1 | 153.1438 | 12.55363 | 12.19916 | **6.39E-30** | Hue 1 |
| RUB:ROS1 | -15.8073 | 8.584783 | -1.84132 | 0.066206 | Hue 1 |
| (Intercept) | 44.86354 | 11.70712 | 3.832158 | **0.000144** | Hue 2 |
| RUB | -4.38909 | 8.775609 | -0.50015 | 0.617207 | Hue 2 |
| ROS1 | 5.966213 | 10.20831 | 0.584447 | 0.559201 | Hue 2 |
| RUB:ROS1 | 13.37783 | 6.980937 | 1.916337 | 0.05593 | Hue 2 |
| (Intercept) | 45.07245 | 15.9481 | 2.826197 | **0.004912** | Hue 3 |
| RUB | -5.62737 | 11.95462 | -0.47073 | 0.638054 | Hue 3 |
| ROS1 | 44.90034 | 13.90633 | 3.22877 | **0.00133** | Hue 3 |
| RUB:ROS1 | 28.94835 | 9.509822 | 3.044047 | **0.002465** | Hue 3 |
| (Intercept) | 37.36295 | 14.20502 | 2.630264 | **0.008812** | Hue 4 |
| RUB | -5.66573 | 10.64802 | -0.53209 | 0.594914 | Hue 4 |
| ROS1 | 75.99549 | 12.38641 | 6.135391 | **1.81E-09** | Hue 4 |
| RUB:ROS1 | 30.53757 | 8.470431 | 3.605197 | **0.000345** | Hue 4 |
| (Intercept) | 23.83616 | 11.92909 | 1.998154 | **0.046277** | Hue 5 |
| RUB | 2.262911 | 8.941996 | 0.253066 | 0.800328 | Hue 5 |
| ROS1 | 133.4256 | 10.40186 | 12.82709 | **1.7E-32** | Hue 5 |
| RUB:ROS1 | 7.261483 | 7.113296 | 1.020832 | 0.30786 | Hue 5 |
| (Intercept) | 23.73608 | 12.04642 | 1.970386 | **0.049382** | Hue 6 |
| RUB | 3.214847 | 9.029942 | 0.356021 | 0.721985 | Hue 6 |
| ROS1 | 132.4046 | 10.50416 | 12.60496 | **1.41E-31** | Hue 6 |
| RUB:ROS1 | 7.46519 | 7.183257 | 1.039249 | 0.299225 | Hue 6 |
